# Study on Liver Sinusoidal Endothelial Cell Fenestrations Based on Cellular Omics-Structure Integration Technology and Its Application in Metabolic Diseases

**DOI:** 10.1101/2025.05.16.653525

**Authors:** Zhuang Wei, Jiji Chen, Maria A. Aronova, Richard D. Leapman

## Abstract

This study developed a new Cellular Omics-Structural Integration (COSI) technology platform to address the limitation of traditional technologies in simultaneously obtaining gene expression profiles and super-resolution cellular structural information at the single-cell level. The platform comprises three core functional modules: (1) a single-cell transcriptomics and super-resolution fluorescence microscopy integration module that enables simultaneous acquisition of gene expression profiles and super-resolution fluorescence images at the single-cell level; (2) an electron microscopy and super-resolution fluorescence microscopy integration module with deep learning resolution enhancement that further gives fluorescence image high resolution features; and (3) a comprehensive analysis module that integrates transcriptomic data with enhanced super-resolution morphological data.

Application of this technology to primary liver sinusoidal endothelial cells successfully achieved efficient matching and analysis of ultrastructural information and gene transcription data at the single-cell level, revealing associations between specific genes and endothelial cell fenestration formation. Through correlation analysis and multivariate statistical methods, we identified specific gene sets associated with fenestration number and average area. Validation in published non-alcoholic steatohepatitis (NASH) and diabetic mouse models demonstrated that these gene sets can effectively assess disease status and drug intervention efficacy, with fenestration number-related gene sets showing significant reduction in NASH and time-dependent changes in response to diabetes treatments.

These findings not only expand our understanding of the mechanisms underlying liver and kidney endothelial cell fenestration formation but also provide novel molecular markers and potential therapeutic targets for early diagnosis and treatment evaluation of metabolic diseases. As a fundamental research tool, COSI technology fills critical gaps in existing spatial omics and cellular biology research, particularly for studying cellular structures lacking specific markers, and demonstrates significant potential for clinical applications in chronic metabolic diseases.

## Introduction

Since the first emergence of single-cell transcriptomics in 2009, single-cell omics has developed rapidly, gradually expanding to multiple dimensions including transcription, genomic variation, epigenetics, proteins, and metabolites (2; 24; 40; 42; 49). Single-cell omics reveals cellular heterogeneity and complex biological processes at the single-cell level, providing unprecedented insights for developmental biology, immunology, and cancer research, while also offering new possibilities for future clinical single-cell detection (19; 22; 41). However, despite emphasizing the “cell” as the fundamental unit, the application of single-cell omics in cell biology, especially in subcellular structural biology, remains relatively limited.

Meanwhile, innovations in microscopy techniques have driven the flourishing development of cell biology over the past decades. Advances in electron microscopy (TEM, SEM, 3D SEM, STEM) and super-resolution fluorescence microscopy (STED, PALM, SIM) have enabled us to observe and analyze subcellular structures with unprecedented spatial resolution (5; 26; 28; 43). However, these technologies are limited by the throughput of targets; even with the latest multi-channel fluorescence technology, the number of simultaneously detectable targets is difficult to exceed a few dozen.

Combining single-cell omics with advanced microscopy techniques promises to overcome these limitations, integrating spatial information with molecular function to form a complete and highly detailed map of cellular structure and function (20). This new integration approach is particularly suitable for studying cell types with special spatial structures, such as fenestrated capillary endothelial cells found in the liver and kidneys (10; 26; 28). These specialized endothelial cells are responsible for material exchange between the liver and kidneys and blood through the formation of fenestrations, playing a crucial role in normal organ function (39).

Studies have shown that early pathological changes in liver and kidney endothelial fenestrations are closely related to the occurrence of chronic liver and kidney diseases. For example, one of the early symptoms of non-alcoholic steatohepatitis (NASH) is the reduction of sinusoidal endothelial cell fenestrations, which seriously affects the exchange of substances between organs and blood, and may further aggravate the pathological state (23; 25). In view of this, we propose that endothelial fenestration pathology may be an important driving factor in the development of liver and kidney-related chronic diseases, therefore exploring its biological mechanisms is crucial for disease prevention and treatment.

However, traditional technologies are limited by the small scale of endothelial fenestrations, making it difficult to reveal the molecular mechanisms of their formation and pathological changes (35). To overcome this technological bottleneck, our laboratory has developed a new technology platform, Cellular Omics-Structural Integration (COSI). As shown in our research results, this system includes three core functional modules: (1) a module integrating single-cell transcriptomics with super-resolution fluorescence microscopy, achieving simultaneous acquisition of genomic expression profiles and super-resolution fluorescence images at the single-cell level; (2) a module combining electron microscopy with super-resolution fluorescence microscopy and deep learning resolution enhancement, further improving fluorescence image features through deep learning technology; (3) a multi-omics data comprehensive analysis module, integrating transcriptomics with enhanced super-resolution morphological data.

Through this new platform, we have achieved efficient matching and analysis of ultrastructural information and gene transcription information at the single-cell level in primary liver sinusoidal endothelial cells, revealing the association between specific genes and endothelial cell fenestration formation. Validation in NASH and diabetic mouse models indicates that these gene sets can be used to assess disease status and drug intervention effects, providing a new perspective for studying the mechanisms of chronic liver and kidney disease development.

Furthermore, our research also provides new possibilities for clinical applications, especially in early diagnosis and treatment evaluation of liver and kidney diseases. The COSI technology platform can be used to screen potential drugs affecting liver and kidney endothelial cell fenestration function, providing new strategies for the prevention and treatment of metabolic diseases. Therefore, this study not only achieves breakthrough progress at the level of basic cell biology but also lays the technological and theoretical foundation for future disease diagnosis and treatment evaluation.

## Results

### System Objectives and Module Design

This research has developed a new data collection and analysis system referred to here as COSI (Cellular Omics-Structural Integration), aimed at achieving two core objectives: (1) simultaneously acquiring whole-genome expression profile data and super-resolution structural information at the same single-cell level; (2) enhancing some high-resolution image features, rather than the resolution of the entire image of super-resolution fluorescence microscopy using deep learning technology to reach electron microscopy levels in certain metrics.

As shown in Figure 1, the system is designed with three functional modules. These three modules can be run separately or together. The first module is the single-cell transcriptome and super-resolution fluorescence microscopy integration module (Figure 1 A-E, blue background), involving precise sorting of individual cells into 384 plates via flow cytometry, followed by fluorescence staining treatment and iSIM (Instant Structured Illumination Microscope) super-resolution imaging (12; 46), while simultaneously performing single-cell mRNA transcriptome sequencing. In this module, individual cells are labeled with an amphipathic molecules CellMask fluorescent dye, super-resolution image matrices are acquired through instant structured illumination microscopy, and mRNA transcriptome sequencing is performed on the same batch of cells to obtain whole-genome expression profile data.

**Figure 1.**
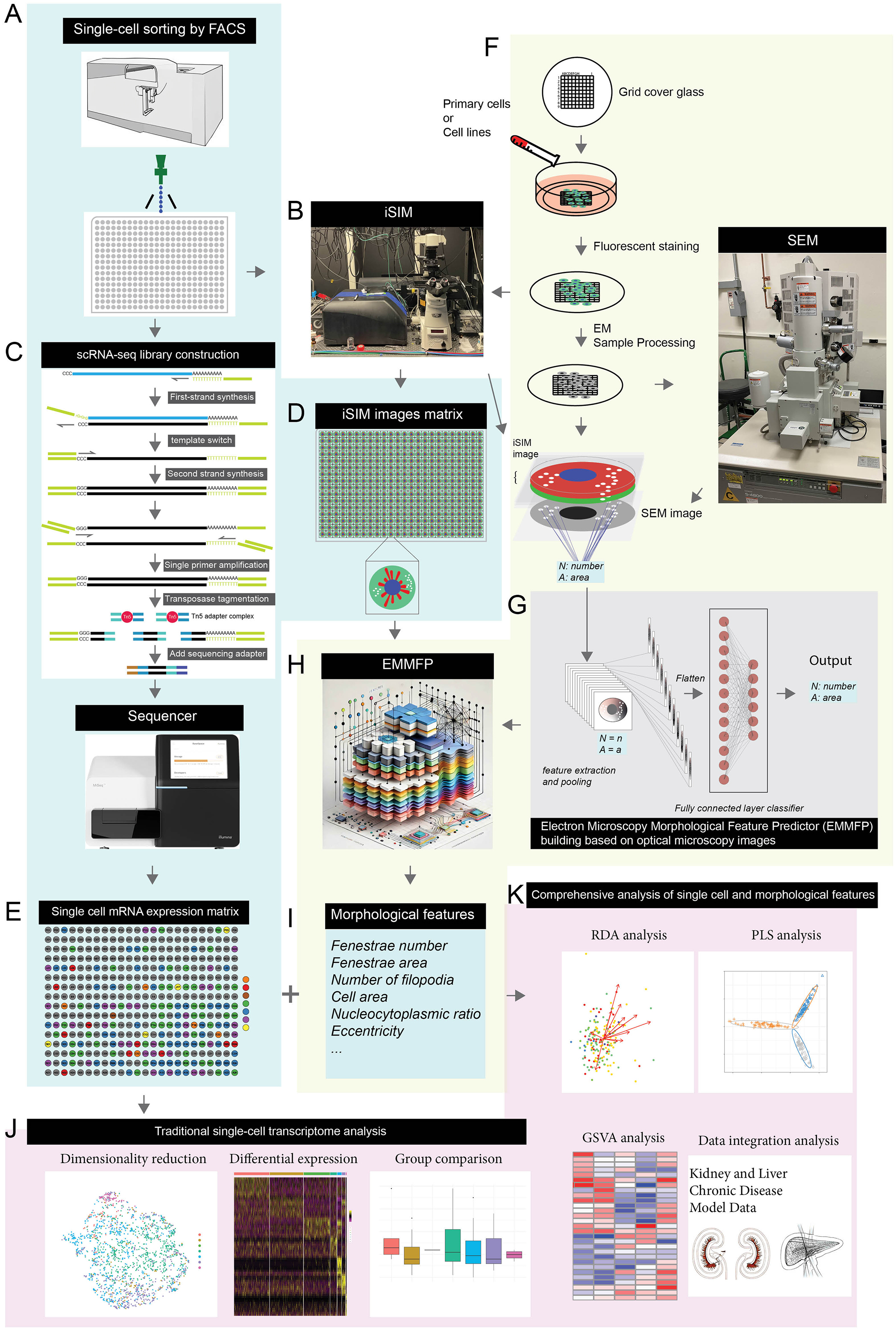
COSI Cellular Omics-Structural Integration): A schematic diagram of the integrated system of single-cell transcriptomics and electron microscopy. This system is designed to simultaneously obtain genome-wide expression profiles and super-resolution structural information at the single-cell level, with further enhancement of imaging resolution through deep learning. (A-E) Single-cell transcriptomics and super-resolution fluorescence microscopy integration module (blue background): (A) Individual cells are precisely sorted into 384 plates via flow cytometry; (B, D) Single cells are labeled with fluorescent dyes and imaged using instant Structured Illumination Microscopy (iSIM) to obtain super-resolution image matrices; (C, E) The same batch of cells is used for mRNA transcriptome sequencing to obtain genome-wide expression profile data. (F-I) Electron microscopy with super-resolution fluorescence microscopy integration and resolution enhancement module (yellow background): (F, G) Electron microscopy is used to obtain cellular ultrastructure information, which is matched with iSIM images; (H, I) Deep learning algorithms enhance super-resolution fluorescence microscopy images to achieve electron microscopy-level spatial index. (J-K) Comprehensive analysis module for transcriptome and super-resolution morphological data, integrating the above data to achieve comprehensive analysis of single-cell multi-omics data, used to study how changes in gene expression affect subcellular structure and function.

The second module is the electron microscopy and super-resolution fluorescence microscopy integration and deep learning resolution enhancement module (Figure 1 F-I, yellow background), which uses electron microscopy to obtain subcellular ultrastructure information and match it with iSIM images, enhancing image resolution through deep learning algorithms. This module mainly addresses the inherent resolution limitations of fluorescence microscopy by matching electron microscopy data with iSIM images through deep learning algorithms to give some super-resolution level image index (image features), enabling them to achieve spatial resolution approaching electron microscopy levels. However, this model still exhibits instances where domains cannot be fully aligned. To address this issue of “domain shift,” we have redesigned the model; details can be found in the supplementary materials (Supplementary Model). However, the subsequent analysis in this paper continues to utilize data derived from the original model. The new model serves solely to illustrate the methodology and architecture.

The third module is the comprehensive analysis module for transcriptome and super-resolution morphological data (Figure 1 J-K), integrating data from the first two modules to achieve comprehensive analysis of single-cell multi-omics data. This module combines gene expression profiles with high-resolution morphological features to study how changes in gene expression affect subcellular structure and function, particularly in the fenestration phenomenon of liver sinusoidal endothelial cells.

### Primary Liver Sinusoidal Endothelial Cells Single-Cell mRNA Sequencing Analysis

Using COSI technology, single-cell mRNA sequencing was performed on primary liver sinusoidal endothelial cells to obtain high-quality transcriptome data (Figure 2 and S1). During the experimental process, we first sorted primary liver sinusoidal endothelial cells into 384-well plates and processed them according to the cell culture and staining protocols described in the materials and methods. Prior to fixation and after fixation and staining, the multi-well plates were placed under an iSIM super-resolution microscope equipped with a high-precision automated stage for cell morphology imaging. The iSIM utilizes two sets of microlens arrays in the optical path to achieve structured illumination of the sample, enabling high-speed structured illumination imaging compared to traditional SIM fluorescence microscopy, making it particularly suitable for high-content super-resolution cellular imaging (12; 46). In our study, iSIM images achieved an XY-axis resolution of 150nm without deconvolution, providing high-dimensional images more suitable for machine learning modeling analysis.

**Figure 2.**
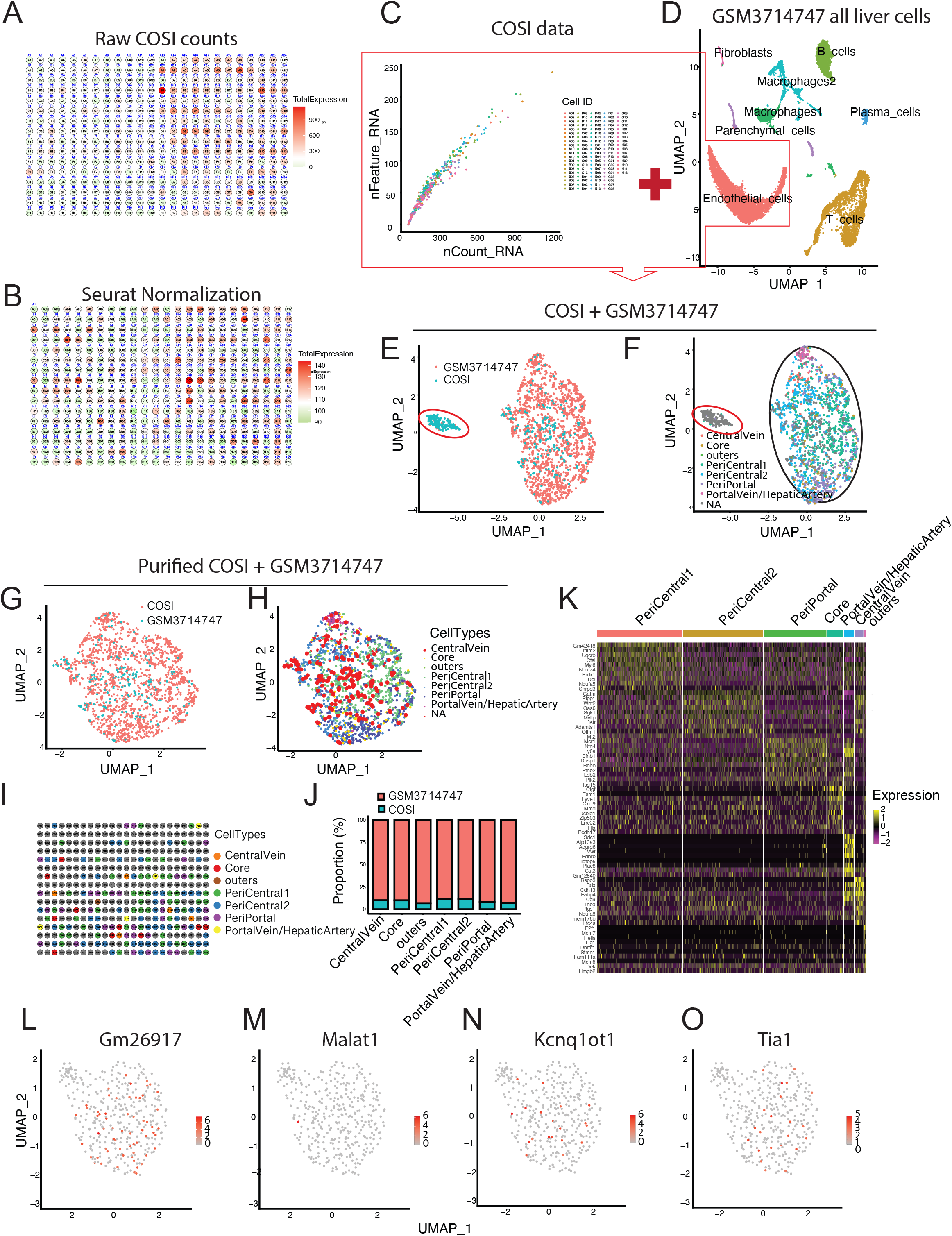
Analysis and validation of primary liver sinusoidal endothelial cell single-cell mRNA sequencing data using COSI technology. A-B. Gene expression matrix diagram in 384-well plate. Panel A shows original non-normalized data, with color gradient representing gene expression levels, showing obvious regional correlation effects; Panel B displays the matrix after Seurat normalization processing, effectively eliminating regional effects. C. Scatter plot showing the relationship between gene counts (nfeature) and sequencing read counts (ncounts) between two sequencing batches. D-F. Data integration and dimensionality reduction analysis. Panel D shows the UMAP plot of the published GSE129516 (GSM3714747) dataset (45), with red clusters representing endothelial cells used for integration validation with COSI data; Panel E shows the integrated UMAP analysis of COSI data (red) and GSE129516 endothelial cell data (blue); Panel F is the integrated UMAP plot with refined endothelial cell subgroup classification (central vein, portal vein/artery, sinusoidal core area, etc.). Black ellipses mark normally integrated endothelial cells. Red circles indicate abnormal clusters that were removed in further analysis. G-H. UMAP plot of purified endothelial cell data. Panel G distinguishes between COSI data (blue) and the GSE129516 dataset; Panel H marks different cell subgroups, particularly highlighting cells from our laboratory’s COSI technology (red, emphasized with large red dots). I. Distribution matrix of endothelial cell types on the 384-well plate. J. Bar chart showing the proportion of integrated endothelial cell data. The X-axis represents cell subgroups, and the Y-axis represents proportions. Blue represents cells from COSI technology, and red represents cells from the GSE129516 dataset. K. Heat map analysis of differentially expressed genes in endothelial cells. The X-axis represents different endothelial cell types, and the Y-axis displays significantly differentially expressed genes, with color gradient indicating expression levels. L-O. Visualization of expression levels of four key differentially expressed genes (*GM26917, Malat1, Kcnq1ot1*, and *Tia1*) on UMAP plots, with color intensity indicating expression levels.

Subsequently, using a modified single-cell RNA extraction and library construction method, the samples proceeded to mRNA library construction. Since mRNA in cells is not actually present in free form but exists bound to various protein and ribosomal RNA complexes, this characteristic protects mRNA from degradation to a large extent, especially under fixed conditions, but also increases the difficulty of releasing mRNA from cells (16; 36). Therefore, we first lysed the cells with a lysis buffer containing proteinase K, with calcium chloride present to enhance proteinase K activity to better release proteins lightly fixed by formaldehyde, exposing mRNA more effectively. Mercaptoethanol provided a reducing environment to further protect mRNA from oxidative damage. Furthermore, we increased the salt concentration of the lysate by adding TCL buffer and sodium chloride to enable RNA binding to magnetic beads for purification in a high-salt environment. In this study, we further employed a method similar to SMART-seq2 for sequencing library construction (29). SMART-seq2 utilizes template switch reactions for first-strand and second-strand synthesis, suitable for full-length sequencing, allowing further mRNA assembly to explore variable splicing and other features.

As shown in Figure 2A-B, the original unnormalized data from the 384-well plate exhibited obvious regional effects, presenting a spatially correlated distribution of gene expression levels. After Seurat normalization processing (37), this regional effect was effectively eliminated, indicating that data preprocessing effectively reduced technical bias. The sequencing data from two batches showed stability in terms of gene number (nfeature) and sequencing reads count (ncounts) (Figure 2C), reflecting that the COSI technology established in this study has good inter-batch consistency and stability.

To verify the biological accuracy of COSI technology data, we performed integrated analysis of our experimental data with the endothelial cell portion from the published PRJNA531644 (GSE129516) dataset (45) (Figure 2D-F, figure S2). After eliminating batch effects using FindIntegrationAnchors and IntegrateData functions, the Uniform Manifold Approximation and Projection (UMAP) dimensionality reduction results showed that our laboratory’s COSI data (blue) and the reference dataset (red) were highly integrated, demonstrating good biological consistency. Refined UMAP analysis classified endothelial cell subgroups as central vein, portal vein/artery, sinusoidal core area and outer area, as well as pericentral1, pericentral2, and periportal regions, with normally integrated cell clusters marked by black ellipses. Abnormal clusters marked by red circles were identified and removed in subsequent analyses to ensure data quality.

The integrated endothelial cell data analysis after removing abnormal clusters (Figure 2G-K, figure S2 C and D) further validated the accuracy of cell type annotation. Figure 2G clearly distinguished COSI data (blue) from the GSE129516 dataset (red), Figure 2H marked different cell subgroups, particularly emphasizing cells obtained using our laboratory’s COSI technology (blue, emphasized with large red dots). Figure 2I displayed the distribution matrix of endothelial cell types on the 384-well plate, reflecting spatial distribution patterns. Figure 2J showed the composition of integrated endothelial cell data through a proportion bar chart, with blue representing cells from COSI technology and red representing cells from the GSE129516 dataset, further confirming that the data obtained by COSI technology showed good concordance with the reference dataset in terms of cell type composition.

Differential expression gene analysis (Figure 2K,figure S2E and Supplementary file 1 - 3) displayed specific marker genes for various endothelial cell subgroups through a heatmap, with the X-axis representing different endothelial cell types and the Y-axis showing significantly differentially expressed genes, with color gradients indicating expression levels. The visualization of expression levels of four key differentially expressed genes (*GM26917, Malat1*, Kcnq1ot1, and *Tia1*) on UMAP plots (Figure 2L-O, figure S2F-H) further demonstrated the molecular characteristics of endothelial cell subgroups, with color intensity indicating expression levels. These results collectively verified that COSI technology could reliably capture the molecular heterogeneity of liver sinusoidal endothelial cells.

### Single-Cell Morphological Features and Gene Expression Correlation Analysis

To explore the association between cellular morphological features and gene expression, we conducted integrated analysis of morphological features and gene expression profiles at the single-cell level on primary liver sinusoidal endothelial cells (Figure 3). Morphological features were obtained through manual annotation and computational models, including fenestration number (Number_of_Holes), average fenestration area (Mean_Hole_Area), nuclear-cytoplasmic ratio (Nuc2Cell), eccentricity (Eccentricity), and aspect ratio (AspectRatio). These parameters were obtained through analysis of high-resolution fluorescence images and matched with transcriptome data from the same cell (Supplementary file 4).

**Figure 3.**
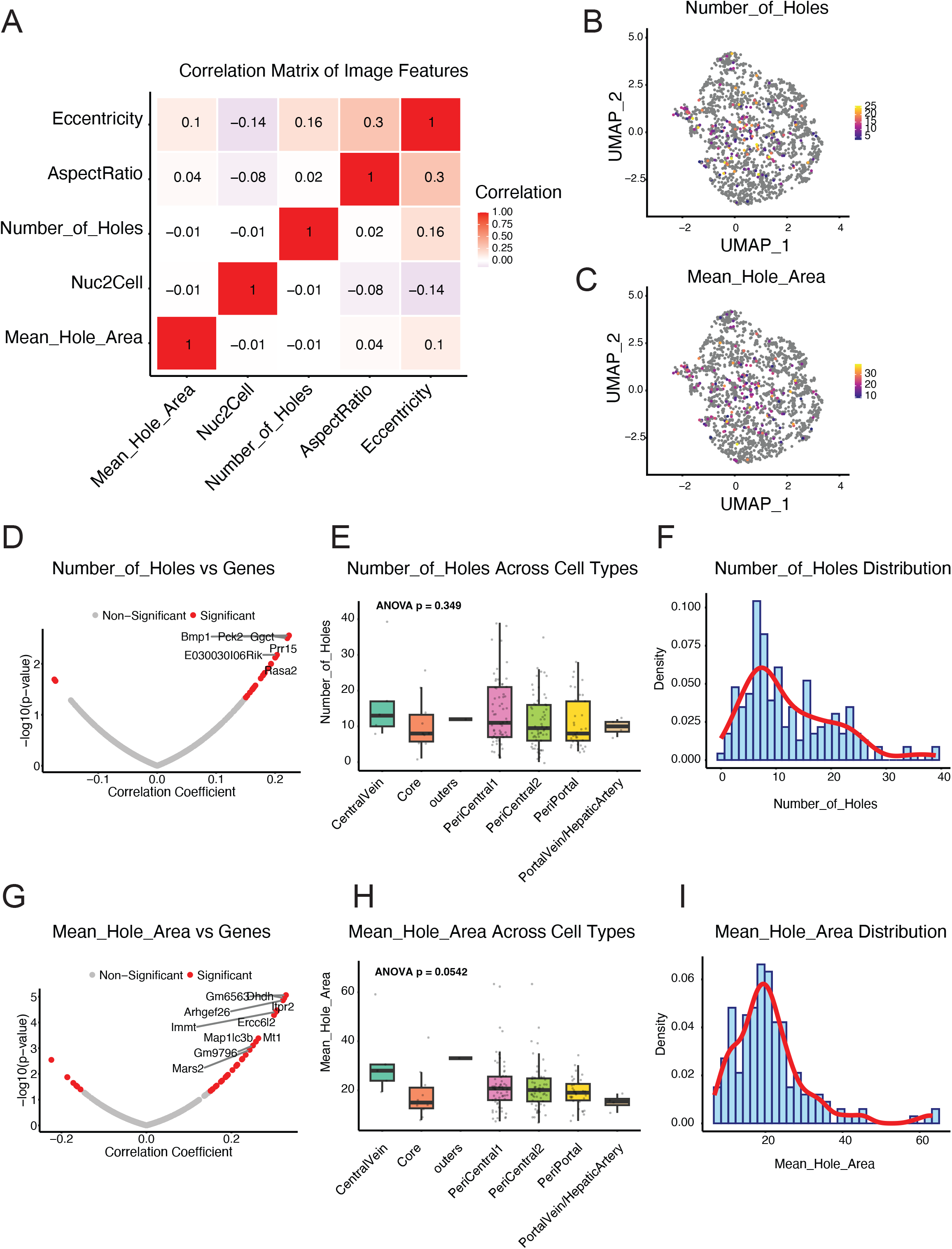
Analysis of the correlation between single-cell mRNA sequencing data and morphological characteristics of primary liver sinusoidal endothelial cells using COSI technology. Multiple morphological parameters of primary liver sinusoidal endothelial cells were obtained through manual annotation and computational models, and then integrated with single-cell mRNA sequencing data for correlation analysis. A. Correlation analysis of liver sinusoidal endothelial cell morphological features. The heat map shows Pearson correlation coefficients among five major morphological features. Significant positive correlations exist between the number of fenestrations and eccentricity (*r*=0.161, *p*=0.032), as well as between aspect ratio and eccentricity (*r*=0.300, *p*<0.001). B-C. Distribution of morphological features in UMAP space. Panel B shows the distribution of Mean_Hole_Area, with colors ranging from yellow (low values) to deep red (high values); Panel C shows the distribution of Number_of_Holes, displaying variability in different regions of LSECs. D and G. Association between image features (mean hole area; number of holes) and gene expression. Volcano plots show the results of correlation analysis between gene expression and image features. The X-axis represents the correlation coefficient (r), and the Y-axis represents - log10 (*p*-value). Red dots indicate highly significantly correlated genes (*p*<0.01), and gray dots indicate non-significant correlations. The top 10 most significantly correlated genes are labeled, including *Gm6563* (*r*=0.328), *Dhdh* (*r*=0.324), *Arhgef26* (*r*=0.321), *Itpr2* (*r*=0.307), and *Immt* (*r*=0.305). E and H. Comparison of morphological features of LSECs in different liver lobule regions. Panel E shows mean hole area, with a gradient trend from the central venous zone to the portal venous zone (*p*=0.054); Panel H shows the number of holes, with differences not reaching statistical significance (*p*=0.349). F and I. Histograms of morphological feature distribution. Panel F shows the distribution of hole numbers; Panel I shows the distribution of mean hole area. Both graphs use blue bars to represent frequency counts and red curves to represent smoothed probability density estimates. The number of holes quantifies the number of cytoplasmic vacuoles or perforations in each cell, related to cell porosity; mean hole area measures the average size of cytoplasmic vacuoles or perforations in each cell, potentially indicating ultrastructural and morphological differences between cell populations.

Correlation analysis among morphological features (Figure 3A, supplementary file 5 and 6) displayed Pearson correlation coefficients between five major morphological features through a heatmap, revealing intrinsic connections in the structural organization of liver sinusoidal endothelial cells (LSECs). The data showed a significant positive correlation between eccentricity and aspect ratio (*r*=0.300, *p*<0.001), consistent with the geometric principles of endothelial cell morphology, indicating that the elongation degree of cell shape is closely related to its asymmetry. Additionally, there was a statistically significant positive correlation between fenestration number and eccentricity (*r*=0.161, *p*=0.032), suggesting that the asymmetry of cell shape might be related to its fenestration formation mechanism, providing a new perspective for understanding the formation of endothelial cell fenestrations.

In UMAP space, the distribution of average fenestration number (Figure 3B) and fenestration area (Figure 3C) presented a relatively random pattern without obvious spatial enrichment trends, with colors ranging from yellow (low values) to deep red (high values) showing the variability of these parameters in LSECs from different regions. This distribution pattern suggests that the formation of endothelial cell fenestrations might be regulated by multiple factors collectively, not limited to linear determination by cell type.

Correlation analysis with gene expression (Figure 3D and G, supplementary file 7 and 8) displayed the correlation analysis results between gene expression and average fenestration area through volcano plots. The X-axis represents the correlation coefficient (*r*), and the Y-axis represents -log10 (*p*-value). Red dots indicate highly significant correlated genes (*p*<0.01), and gray dots indicate non-significant correlations. The top 10 most significantly correlated genes are marked, including *Gm6563* (*r*=0.328), *Dhdh* (*r*=0.324), *Arhgef26* (r=0.321), *Itpr2* (*r*=0.307), and *Immt (r*=0.305). These genes may participate in regulating the formation and maintenance of liver sinusoidal endothelial cell fenestration structures, providing important clues for subsequent functional research.

Analysis based on liver lobule regional characteristics (Figure 3E and H) showed that LSECs from different regions exhibited certain region-specific characteristics in morphological features. From the central vein region to the portal vein region, the average fenestration area showed a gradient change trend along the direction of the hepatic lobular sinusoidal endothelial cells from the portal vein to the central vein. (*p*=0.054), indicating that different anatomical locations of the liver lobule might affect the size of endothelial cell fenestrations. In contrast, regional differences in fenestration number did not reach statistical significance (*p*=0.349), suggesting that fenestration number might be influenced by more complex regulatory mechanisms.

The distribution histograms of liver sinusoidal endothelial cell fenestration number and area (Figure 3F and I, supplementary file 4) further quantified the statistical characteristics of these morphological features. The fenestration number (Figure 3F) averaged 12.41±8.20, with a distribution range of 0-39; the average fenestration area (Figure 3I) was 21.20±9.92μm^2^, with a median of 19.61μm^2^, and showed a strong positive skew distribution. These data reflect the heterogeneity of LSEC fenestration structures and provide important baseline data for studying endothelial cell fenestration function. Fenestration number quantifies the number of cytoplasmic vacuoles or perforations in each cell, related to cell porosity; average fenestration area measures the average size of cytoplasmic vacuoles or perforations in each cell, potentially indicating ultrastructural and morphological differences between cell populations.

### Multivariate Analysis of Morphological Features and Gene Expression

To comprehensively understand the complex relationship between morphological features and gene expression, we employed various multivariate statistical analysis methods. First, we applied Redundancy Analysis (RDA) to explore the association between cellular morphological features and gene expression patterns (Figure 4A - C and supplementary file 9 - 14). RDA is a constrained ordination method that can directly analyze the influence of morphological features (environmental variables) on gene expression (response variables).

**Figure 4.**
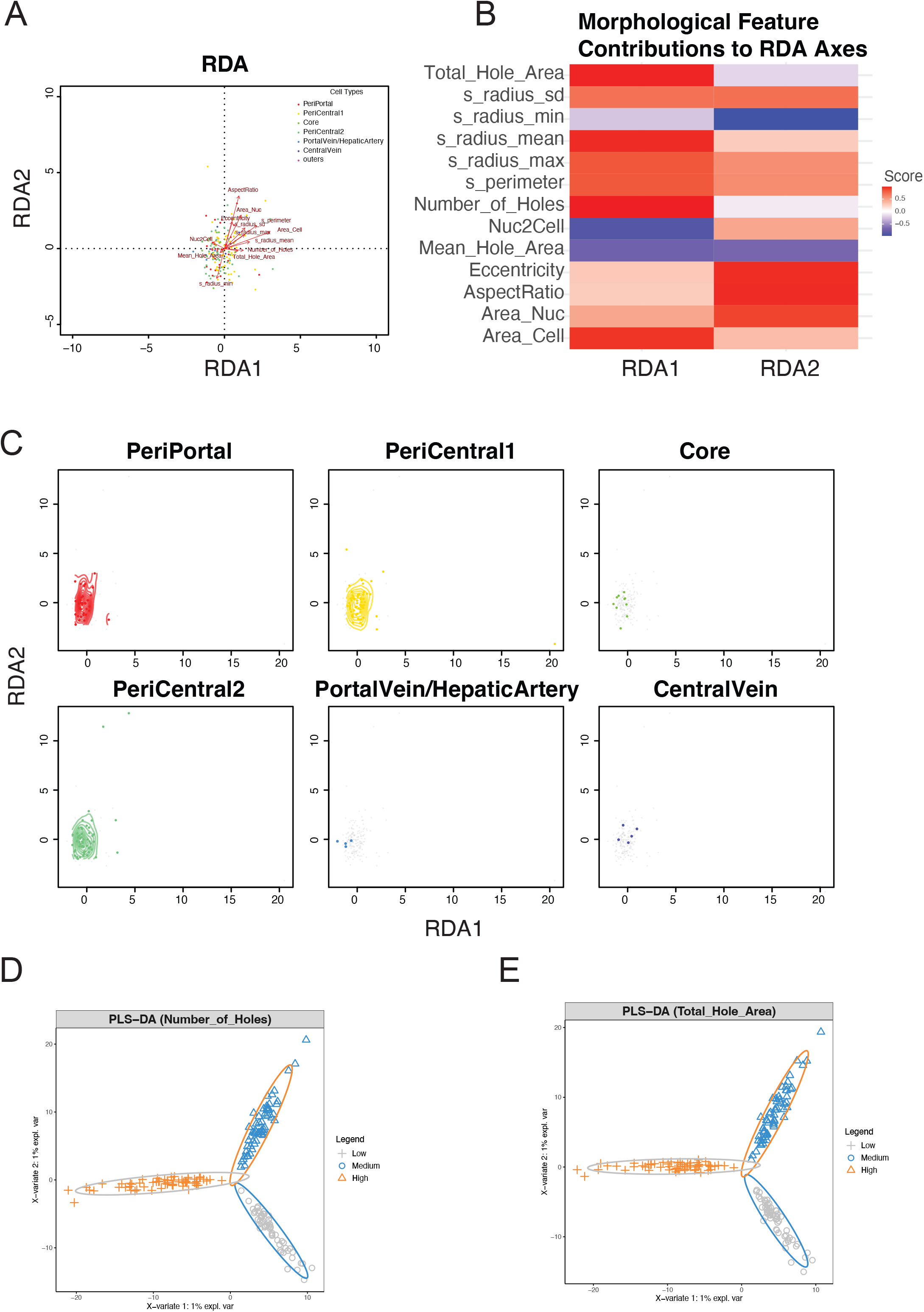
Integrated analysis of morphological features and gene expression. A: Basic RDA plot Redundancy Analysis (RDA) results showing the relationship between gene expression and cell morphological features. Each point represents a cell, with colors distinguishing different cell types. Red arrows indicate the direction and strength of morphological feature influences, arrow length represents the degree to which features explain gene expression variation, and arrow direction shows the correlation of features with RDA axes. The X-axis (RDA1) and Y-axis (RDA2) explain the largest and second largest proportions of variation, respectively. B: Heat map of morphological feature contributions Displays the contribution of various morphological features to the RDA1 and RDA2 axes. Colors represent the direction and strength of correlation: blue for negative correlation, white for no correlation, and red for positive correlation, with deeper colors indicating stronger correlations. C: Individual cell type distribution plot Multi-panel plot showing the distribution density of each cell type in RDA space. Gray dots represent the background distribution of all cells, colored dots represent specific cell types, and colored contour lines (when samples are sufficient) show the density distribution of cell types. X/Y axes are RDA1 and RDA2, respectively. D-E: PLS-DA analysis of cell parameters Partial Least Squares Discriminant Analysis (PLS-DA) showing the distribution of cells classified based on the number of holes (D) and mean hole area (E) parameters, including low (blue), medium (green), and high (red) groups. Ellipses represent the 95% confidence interval for each group. X and Y axes represent the first and second principal components, respectively.

In the basic RDA plot (Figure 4A), each point represents a cell, with colors distinguishing different cell types. Red arrows indicate the direction and strength of morphological feature influences, arrow length represents the extent to which features explain gene expression variation, and arrow direction shows the correlation of features with RDA axes. Cell types exhibited certain separation patterns in RDA space, with morphological features contributing differently to gene expression variation in different directions. The X-axis (RDA1) and Y-axis (RDA2) explain the largest and second-largest proportions of variation, respectively. Although the overall RDA model showed statistical significance (*p*=0.004), these morphological features could only explain 7.93% of the total variation in gene expression, indicating that gene expression is influenced by multiple factors, with cell morphology being just one of them.

The morphological feature contribution heatmap (Figure 4B) displayed the contributions of various morphological features to the RDA1 and RDA2 axes. Colors indicate the direction and strength of correlation: blue for negative correlation, white for no correlation, and red for positive correlation, with deeper colors indicating stronger correlations. More image features include cell area (Area_Cell), average radius (s_radius_mean), nuclear-cytoplasmic ratio (Nuc2Cell), and nuclear area (Area_Nuc) had significant impacts on gene expression, while aspect ratio (AspectRatio) showed the strongest correlation on the RDA2 axis.

The distribution of various cell types in RDA space (Figure 4C) displayed the density distribution of each cell type in RDA space through multi-panel plots. Gray points represent the background distribution of all cells, colored points represent specific cell types, and colored contour lines show the density distribution of cell types. The X/Y axes are RDA1 and RDA2, respectively. This analysis further confirmed the specificity of different cell types in terms of morphology-gene expression relationships, suggesting that the association between morphological features and gene expression might be cell type-specific.

Additionally, To evaluate the correlation between morphological features and gene expression, we conducted Partial Least Squares Discriminant Analysis (PLS-DA)(34) on cell classifications based on pore count (Figure 4D) and pore area (Figure 4E). Results indicated these parameters could distinguish different cell populations to some extent. The analysis using image features involved dividing cells into three equal groups (low, medium, and high image features). We identified specific gene sets (Top 30 genes) associated with various morphological features (Supplementary file 15 - 28). These morphology-specific gene sets not only support the value of morphological classification in cell type identification but also provide important clues for understanding the molecular basis of cellular morphological diversity, which can be utilized for subsequent functional enrichment and pathway analysis.

### Functional Analysis of Liver Endothelial Cell Fenestration-Related Gene Sets and Pathological Model Validation

Based on the PLS-DA method, we constructed specific gene sets associated with five key morphological features (Mean_Hole_Area, Nuc2Cell, Number_of_Holes, AspectRatio, and Eccentricity). To further investigate the relationship between these gene sets and previously reported signaling pathways and biological processes, we performed Gene Set Variation Analysis (GSVA) and correlation analysis (13). GSVA is a method that transforms gene expression data into gene set activity scores, enabling research to extend from single-gene level to functional pathway level. We utilized gene sets from the Molecular Signatures Database (MSigDB), which contains functionally annotated gene sets from various sources, covering biological pathways, transcription factor regulatory networks, and other multi-level functional classifications (6; 38). Subsequently, we calculated the correlation between the GSVA scores of all mouse gene sets in MSigDB and morphological features. Results showed revealed distinct molecular signatures associated with cellular morphological features (Supplementary file 29 - 40). For Mean_Hole_Area, the strongest negative correlation was observed with GSE12198_CTRL_VS_LOW_IL2_STIM_NK_CELL_DN (*r* = -0.317, *p* = 1.76e^-5^), suggesting that genes downregulated in NK cells under low IL-2 stimulation may influence hole area reduction (Supplementary file 30, 32 and 34). Several Golgi-related pathways also showed significant negative correlations, including vesicle transport regulation and cisterna membrane components (both *r* = -0.277, *p* = 2.04e^-4^), indicating potential involvement of secretory pathways in controlling this morphological feature. Conversely, ACEVEDO_LIVER_CANCER_UP showed the strongest positive correlation (*r* = 0.271, *p* = 2.81e^-4^), linking liver cancer-associated genes with increased hole areas. For Number_of_Holes, multiple pathways shared identical strong negative correlations (*r* = -0.323, *p* = 1.17e^-5^), notably BIOCARTA_AKAPCENTROSOME_PATHWAY and several morphology-related gene sets, suggesting centrosome regulation may influence hole formation. Positive correlations were led by NF1_Q6 (*r* = 0.316, *p* = 1.80e^-5^) and POOLA_INVASIVE_BREAST_CANCER_UP (r = 0.313, *p* = 2.21e^-5^), potentially linking NF1-regulated genes and invasive cancer signatures to increased hole formation (Supplementary file 31, 33 and 36). These findings highlight the molecular underpinnings of cellular morphological features and suggest potential regulatory mechanisms connecting gene expression patterns to structural changes observable in imaging analyses, more correlation analyses between biological processes and image features are provided in the supplementary figures (Supplementary file 37 - 40).

Furthermore, we analyzed the distribution of significantly correlated gene sets in image features (Figure 5A). Green represents the proportion of positively correlated gene sets (*p* < 0.01), and red represents the proportion of negatively correlated gene sets. Bar charts are sorted from high to low by the proportion of positively correlated gene sets. The results showed significant heterogeneity in the distribution of positive and negative correlations across different image features. Notably, features such as eccentricity (Eccentricity), nuclear-cytoplasmic ratio (Nuc2Cell), fenestration number (Number of Holes), and average fenestration area (Mean hole area) primarily showed positive correlations, while aspect ratio (AspectRatio) displayed a higher proportion of negative correlations. This differential distribution pattern might reflect different biological processes and regulatory mechanisms behind various cellular phenotypes. Figure 5B summarized the biological functions corresponding to MSigDB gene sets associated with five key morphological image features, providing important clues for understanding the molecular basis of these morphological features.

**Figure 5.**
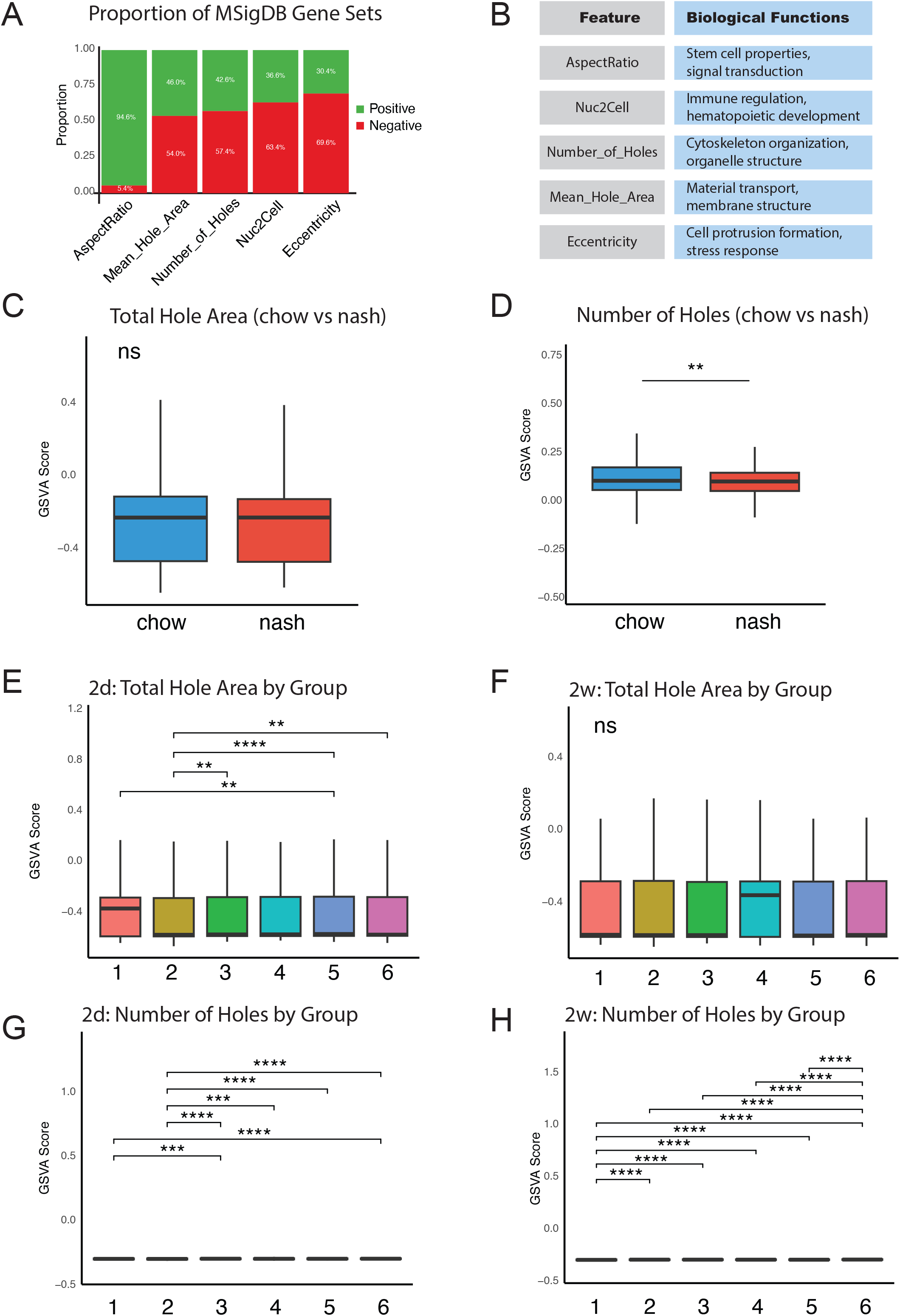
Significance of COSI-discovered endothelial cell fenestration-related gene sets in liver and kidney chronic disease models. A. Distribution of significantly correlated MSigDB gene sets (6; 38) in image features (*p* < 0.01). Green represents the proportion of positively correlated gene sets, and red represents the proportion of negatively correlated gene sets. Bar charts are sorted from high to low by the proportion of positively correlated gene sets. B. Functional summary of MSigDB gene sets associated with five key morphological image features with Keyword Frequency Analysis. C-D. Box plots comparing GSVA scores between chow (control) and nash groups of (GSE129516) dataset (. Panel C shows the Total_Hole_Area gene set (blue), and Panel D shows the Number_of_Holes gene set (red). Wilcoxon rank-sum test indicates that the Number_of_Holes gene set shows significant differences between the two groups (*p*=0.001312), while the Total_Hole_Area gene set differences are not significant (*p*=0.2772). ** *p*<0.01, ns: not significant. E-H. Box plots of GSVA scores for different treatment groups at different time points in Diabetic kidney disease model data (GSE184652)(44). Panels E and G show the Total_Hole_Area gene set, and panels F and H show the Number_of_Holes gene set. E-F are for the 2d time point, and G-H are for the 2w time point. Kruskal-Wallis test shows: at the 2d time point, both gene sets have significant differences (Total_Hole_Area: *p*=0.0004215; Number_of_Holes: *p*=2.34×10^-12); at the 2w time point, the Total_Hole_Area gene set shows no significant difference (*p*=0.2282), while the Number_of_Holes gene set shows highly significant differences (*p*=6.238×10^-29). The X-axis represents groups: 1.db/db+AAV+PBS 2.db/db+AAV+ACEi 3.db/db+AAV+Rosi 4.db/db+AAV+ACEi+Rosi 5.db/db+AAV+SGLT2i 6.db/db+AAV+ACEi+SGLT2i, and the Y-axis represents GSVA scores. \**p*<0.05,\*\**p*<0.01, \*\*\**p*<0.001,\*\*\*\**p*<0.0001.

To further investigate the role of our identified gene sets in chronic liver and kidney diseases, we reanalyzed data from a liver NASH mouse model (GSE129516)(45) and a diabetic mouse model dataset (GSE184652)(44).

In the analysis of liver NASH mouse model (GSE129516) data (Figure 5C-D, supplementary file 41 - 45), we compared the GSVA scores of two key gene sets in endothelial cells between the normal diet group (chow as control) and the NASH group. Wilcoxon rank-sum test showed that the Number_of_Holes gene set exhibited significant differences between the two groups (*p*=0.001312), while the Total_Hole_Area gene set did not show significant differences (*p*=0.2772). This suggests that non-alcoholic steatohepatitis might primarily affect vascular function by reducing the expression of genes related to fenestration number rather than changing the total fenestration area, providing a new perspective for understanding liver sinusoidal endothelial cell dysfunction in NASH.

For the diabetic mouse model dataset (GSE184652), we analyzed the effects of different therapeutic interventions on the expression of endothelial cell fenestration-related genes at early (2d) and late (2w) time points (Figure 5E-H, supplementary file 46 - 49). At the 2d time point, Kruskal-Wallis tests showed significant differences among different treatment groups in both Total_Hole_Area (*p*=0.0004215) and Number_of_Holes (*p*=2.34×10^-12^) gene sets. At the 2w time point, the group differences in the Total_Hole_Area gene set weakened (*p*=0.2282), while the differences in the Number_of_Holes gene set became more pronounced (*p*=6.238×10^-29^). This temporal dynamic change revealed that different therapeutic interventions had different short-term and long-term effects on endothelial cell fenestration function.

Further analysis of expression changes in different treatment groups at two time points (Supplementary file 46 - 49) revealed that various interventions had significant time-dependent regulatory effects on fenestration-related genes. The ACEi (Angiotensin-Converting Enzyme inhibitor) treatment group exhibited the most significant time-dependent changes, while Rosi (Rosiglitazone) related treatments showed relatively stable expression between the two time points. SGLT2i (Sodium-Glucose Co-Transporter 2 inhibitor) monotherapy primarily affected the Total_Hole_Area gene set, while ACEi+SGLT2i combination therapy mainly affected the Number_of_Holes gene set. These results suggest that different drugs might improve the fenestration function of renal endothelial cells under diabetic conditions through different molecular mechanisms, providing important references for optimizing clinical treatment strategies.

Combining these findings, we confirmed that COSI technology could successfully integrate single-cell transcriptome and morphological data, revealing the molecular mechanisms of endothelial cell fenestration formation, and through validation in NASH and diabetes models, demonstrated the important role of fenestration-related genes in disease occurrence and treatment response. These results not only expand our understanding of the mechanisms of liver and kidney endothelial cell fenestration formation but also provide new molecular markers and potential intervention targets for early diagnosis and treatment evaluation of chronic metabolic diseases.

## Discussion

Our developed COSI (Cellular Omics-Structural Integration) technology fills a critical gap in current spatial omics and cellular biology research. Unlike existing technologies, COSI is specifically designed for organelle biology research, integrating high-content super-resolution fluorescence microscopy with single-cell mRNA sequencing (1; 8; 9; 18; 31; 32). While existing spatial transcriptomics technologies (such as Visium, Slide-seq, Stereo-seq, etc.) can provide spatial molecular information at the tissue level (21; 30; 47), they have inherent limitations when addressing organelle biology questions, primarily due to insufficient resolution to precisely capture organelle structures, reliance on RNA rather than protein analysis (making it difficult to accurately reflect organelle functional states), and their typical unsuitability for adherent cultured cells, which complicates modeling cell morphology under physiological conditions (11).

In contrast, COSI technology is based on super-resolution imaging, using iSIM (instant Structured Illumination Microscopy) technology to achieve 150nm resolution on the XY axis, obtaining high-dimensional image information without deconvolution processing (12; 46). This far exceeds traditional spatial omics technologies, providing a foundation for detailed organelle structure analysis. Through a multi-well plate adherent cell culture system, COSI mimics the physiological state of cells attached to the basement membrane or extracellular matrix, creating a more realistic in vivo environment. It uses flow cytometry to precisely separate individual cells into 384-well plates (expandable to 1536-well plates), achieving true single-cell level analysis. COSI also introduces the concept of “microscopic image dimensions,” emphasizing that images obtained through antibody staining or dye labeling contain rich texture and morphological features, providing multi-dimensional data sources for machine learning analysis.

COSI technology overcomes numerous technical challenges in practice. Through systematic screening, we identified microplate materials that are heat-resistant and compatible with super-resolution microscopy, resolving the contradiction between temperature sensitivity and optical transparency. For the binding state of mRNA and protein complexes in fixed cells, we designed a special lysis buffer containing proteinase K (with activity enhanced by calcium chloride), β-mercaptoethanol (providing a reducing environment), and other components to effectively release and protect mRNA. For flat-bottom multi-well plates, we improved PCR instrument design to enhance temperature uniformity and adopted a SMART-seq2-like TSO reaction for library construction (40), achieving full-length mRNA sequencing. Through process optimization, we achieved single-cell library construction on 384-well plates, calculating that four 1536-well plates could reach the throughput of one 10X single-cell sequencing sample, laying the foundation for large-scale applications. Although the current version has limited gene capture efficiency (raw data reliably captures only hundreds of genes), we have constructed a ST RT-seq-like (Single-cell Tagged Reverse Transcription sequencing) adapter probe system aimed at achieving 3’ end capture sequencing when expanded to 1536-well plates, expected to increase gene coverage to genome-wide levels (27).

To validate COSI technology’s practicality, we selected fenestrae—a subcellular structure difficult to study using traditional methods—as our research subject. Fenestrae are pore structures less than 100nm in diameter on endothelial cells, playing key roles in liver and kidney physiological and pathological processes, but lacking specific markers and involving complex multi-gene network regulation in their biogenesis (10; 14; 23; 25; 35; 39; 48). After analyzing mouse liver sinusoidal endothelial cells using COSI technology, we successfully detected known liver sinusoidal endothelial cell related genes (such as *Tia1*)(15), validating the method’s reliability. More importantly, we found that bone morphogenetic protein 1 (*Bmp1*) significantly correlates with fenestrae number, while dihydrodiol dehydrogenase (*Dhdh*)(3) correlates with average fenestrae area. *Bmp1*, as a metalloproteinase with activity in cleaving collagen precursors, is consistent with extracellular matrix remodeling during fenestrae formation (7). Through GSVA analysis, we found that liver cancer upregulated gene sets and downregulated gene sets after drug treatment significantly positively correlate with fenestrae area, suggesting potential connections between fenestrae changes and metabolic diseases. These findings were further validated in disease models: in NASH mouse models, fenestrae number-related gene sets were significantly suppressed, consistent with previous reports; in diabetic mouse kidneys, fenestrae-related genes showed significant differences after treatment with ACEi, rosiglitazone, SGLT2i, and other drugs, especially noticeable after two weeks of treatment.

Based on integrated data obtained through COSI technology, our results supports a growing perspective that fenestrae play a “driver” rather than simply a “phenomenon” role in chronic liver diseases (25). In the NASH disease process, initial hepatocyte damage and regeneration lead to liver fibrosis and vascular network reconstruction, then inducing portal hypertension, which further promotes defenestration of sinusoidal endothelial cells (4). Defenestration may in turn exacerbate portal pressure, forming a vicious cycle ultimately leading to the development of NASH, cirrhosis, or liver cancer (17). In diabetic nephropathy drug intervention studies (44), changes in fenestrae-related gene expression patterns suggest different drugs may affect fenestrae structure through different mechanisms: ACEi, as an angiotensin-converting enzyme inhibitor, directly acts on endothelial cells, with its effect on fenestrae formation possibly related to vascular tension regulation (33); rosiglitazone, as a PPARγ agonist, may indirectly regulate fenestrae formation by affecting intracellular lipid metabolism and thereby changing membrane lipid composition. It has been reported that lanifibranor, a pan-peroxisome proliferator-activated receptor (PPAR) agonist, can improve the formation of fenestrae in liver sinusoidal endothelial cells, indicating that our hypothesis is likely to be valid. However, since the PPAR nuclear receptor family has many members and complex functions, its mechanism on fenestrae formation needs further study; SGLT2i, although primarily acting on proximal tubule cells, may indirectly affect fenestrae formation through inducing endothelial cells’ perception and adaptive response to changes in glucose metabolism. These mechanistic hypotheses require further experimental validation but have provided new research directions for understanding fenestrae regulatory networks.

COSI technology has broad application prospects. As a basic research tool, COSI provides a new method for organelle biology research that integrates morphological and molecular characteristics, particularly suitable for studying cellular structures lacking specific markers. Clinically, through standardized upgrades, COSI technology can be developed into a clinical testing tool for precise diagnosis and prognosis assessment of non-alcoholic fatty liver disease, diabetic nephropathy, hypertensive nephropathy, cancer, or other chronic diseases. For example, it could predict disease development trends or drug responses by detecting fenestrae-related gene expression patterns in patient tissue samples. In drug development, pharmaceutical companies can connect COSI technology with existing high-content screening platforms to simultaneously evaluate the effects of drug candidates on organelle morphology and gene expression, improving drug targeting and reducing toxic side effects. Additionally, integrating multi-omics information helps analyze patient disease subtypes, providing a basis for personalized treatment selection, such as predicting diabetic patients’ responses to different hypoglycemic drugs. In conclusion, COSI technology opens new pathways for cellular biology research, especially in studying organelle function, by bridging the gap between subcellular structure and single-cell transcriptomics. With the rapid development of artificial intelligence deep learning and biological computer simulation, I believe data at this level can make critical contributions to the complete computational prediction of development processes, from DNA to the functional development of entire organisms.

## Materials and Methods

Specific materials and methods are described in the supplementary material.

## Supporting information

Supplemental Materials and Methods and images

## Acknowledgments

This work was supported by the intramural program of the National Institute of Biomedical Imaging and Bioengineering, National Institutes of Health. We sincerely thank Ferenc Livak and his colleagues from the Flow Cytometry Core in National Cancer Institute for support in flow cytometry. We sincerely thank The NIH Intramural Sequencing Center (NISC) for sequencing technology support.

## Author Contributions

R.D. Leapman and Z. Wei conceptualized this research project and developed the experimental and analytical plans together with other collaborators. Z. Wei completed the majority of experiments and bioinformatics analyses. J. Chen developed the automated iSIM (instantaneous Structured Illumination Microscopy) imaging workflow, while Z. Wei and J. Chen jointly performed the iSIM imaging. Z. Wei and M. Aronova carried out the scanning electron microscopy experiments. The manuscript was written by Z. Wei and R.D. Leapman.

## Additional information

Supplemental information is appended to this paper.

Competing financial interests: There are no competing financial interests.

