## Supplemental Materials and Methods and images for "Study on Liver Sinusoidal Endothelial Cell Fenestrations Based on Cellular Omics-Structure Integration Technology and Its Application in Metabolic Diseases"

#### Supplementary Materials and Methods

##### Cell Culture and Immunofluorescence Staining

##### Cell Culture and Fixation

This study used 384-well plates to ensure solutions completely cover the cells, improving fixation and staining effects. The specific steps are as follows:

1. Seed cells in 384-well plates and culture for 3.5 hours.
2. Add 2 µl of RPMI 1640 medium containing 3X CellMask (final concentration 1X) to each well. The total volume per well is 7 µl; in actual use, add 5 µl of serum-free RPMI 1640 containing 1X CellMask.
3. Incubate for 30 minutes.
4. Wash twice with PBS, then add 5 µl of 4% paraformaldehyde (PFA) and fix for 15 minutes.
5. Wash three times with PBS.
6. Add 2 µl of staining solution (10 ml staining solution contains: 2 µl 1X DAPI, 25 µl 1X F-actin dye, 10 µl 1X CellMask). In actual experiments, add 5 µl of staining solution per well.
7. Fix with 4% PFA for 15 minutes.
8. Examine staining results using a fluorescence microscope to confirm successful staining.

##### Scanning Electron Microscopy Sample Preparation

This study used Grid slides to ensure that the same cell coordinates can be located in both iSIM microscopy and scanning electron microscopy for correlative optical and electron microscopy:

1. Seed cells onto Grid slides and culture for 3.5 hours.
2. Add 2000µl of RPMI 1640 medium containing 3X CellMask to each well (final concentration 1X).
3. Incubate for 30 minutes.
4. Wash twice with PBS, then add 500µl of 4% paraformaldehyde (PFA) and fix for 15 minutes.
5. Wash three times with PBS.
6. Add 2000µl of staining solution (10ml staining solution contains: 2µl of 1X DAPI, 25µl of 1X F-actin dye, 10µl of 1X CellMask).
7. Post-fix with 4% PFA for 15 minutes.
8. Perform iSIM microscopy imaging.
9. Fix the sample with 2.5% glutaraldehyde for 15 minutes after iSIM imaging.
10. After water rinse, fix the sample with 1% osmium tetroxide for 30 minutes.
11. After water rinse, treat with 1% tannic acid.
12. After water rinse, fix the sample with 1% osmium tetroxide for 30 minutes.
13. After water rinse, stain with 2% uranyl acetate overnight at 4°C.

14. Scanning electron microscopy imaging:
    a. Equipment: Hitachi S-4800 scanning electron microscope (Serial number: HI-9034-0006)
    b. Imaging parameters:
        i. Accelerating voltage: 10 kV
        ii. Emission current: 9800 nA
        iii. Working distance: 10.8 mm
        iv. Magnification: 3000×
        v. Lens mode: Normal
        vi. Scan speed: Capture_FAST2(64)
    c. Image acquisition:
        i. Image size: 2560×1920 pixels
        ii. Pixel size: 16.54 nm
        iii. Signal type: SE(M) (Secondary Electron)

**Single-cell RNA Sequencing**

**Single-cell RNA Extraction and Library Construction**

Refer to the SMART-seq2 method (6; 8). The specific steps are as follows:

**Cell Lysis**

1. Prepare lysis buffer: 0.3% 2-mercaptoethanol, 0.1% Triton X-100, 200 mM Tris-HCl (pH 8.0), 1 mM $CaCl_2$, Proteinase K (1:100 dilution).
2. Add 3 µl of lysis buffer to each well.
3. Incubate at 54°C for 1 hour (lid temperature 60°C).
4. Incubate at 98°C for 10 minutes (lid temperature 110°C) to heat-denature Proteinase K.

**mRNA Enrichment**

1. Prepare high-salt magnetic bead mixture: 7% beads, 47% TCL buffer, 47% 5M NaCl.
2. Add 5 µl of the mixture to each well.
3. Wash twice with 80% ethanol.

**Reverse Transcription**

1. Add primer mixture: oligo dT-VM (0.06 µM), dNTP (1.2 µl), $H_2O$ (1.74 µl), total 3 µl/well.
2. Incubate at 70°C for 5 minutes.
3. Add reverse transcriptase mixture: TS buffer (1.5 µl), TSO primer (0.225 µl), TS enzyme (0.6 µl), $H_2O$ (0.625 µl), total 3 µl/well.
4. Reaction conditions: 42°C for 90 minutes (lid 45°C), 85°C for 5 minutes (lid 90°C), hold at 4°C.

**Second Strand Synthesis and Pre-amplification**

1. Transfer the reverse transcription products to 384-well plates.
2. Reaction system per tube: First strand product (5 μl), KAPA premix 2X (6.25 μl), ISPCR primer 100 μM (0.0125 μl), $H_2O$ (1.2375 μl), total volume 12.5 μl.
3. PCR program: 98°C for 45 seconds; (98°C for 10 seconds, 66°C for 15 seconds, 72°C for 3 minutes) × 20 cycles; 72°C for 5 minutes.
4. Add 12.5 μl DNA magnetic beads to the PCR product.
5. Wash twice with 80% ethanol, dissolve in 8 μl water.

**Sample Barcoding**

Barcoding method was based on the instructions of barcoding kit of seqWell:

1. Add to each well: 2 μl sample barcode reagent, 2 μl DNA, 2 μl encoding buffer.
2. Incubate at 55°C for 15 minutes, hold at 25°C.
3. Add 3 μl X termination solution.
4. Incubate at 68°C for 10 minutes, hold at 25°C.
5. Combine all corresponding wells into one tube.
6. Measure liquid volume with a pipette, add equal volume of MAGwise magnetic beads.
7. Wash twice with 80% ethanol.
8. Add 41 μl of 10 mM Tris-HCl (pH 8.0) to elute the library, transfer 39 μl to a new tube, add 5 μl of the corresponding pooling barcode reagent and 22 μl encoding buffer.
9. Incubate at 55°C for 15 minutes, hold at 25°C.
10. Add 33 μl X termination solution.
11. Incubate at 68°C for 10 minutes, hold at 25°C.

**Final Library Amplification and Purification**

1. Mix the four pooled samples, add an equal volume of MAGwise magnetic beads from the kit, wash twice with 80% ethanol.
2. Elute with 25 μl Tris-HCl.
3. Take 23 μl of eluted DNA, add 27 μl KAPA HiFi PCR premix to a total volume of 64 μl.
4. PCR program: 72°C for 10 minutes; 98°C for 45 seconds; (98°C for 10 seconds, 66°C for 15 seconds, 72°C for 3 minutes) × 12 cycles; 72°C for 3 minutes.
5. Take half of the product, purify with MAGwise, and perform PCR again.
6. After electrophoresis verification, send the product for sequencing.

**RNA Sequencing Data Processing**

**Read Mapping and Alignment**

FastQC was used for preliminary quality control of raw sequencing reads, evaluating sequence quality, GC content, and adapter contamination. For each sample, paired-end reads (R1 and R2) were aligned to the mouse reference genome (GRCm38) using HISAT2 with default parameters and the '--dta' flag to optimize output for transcript assembly (3). The alignment process used four processing threads to ensure computational efficiency.

Although HISAT2 was the primary alignment tool, STAR was also implemented as an alternative aligner to compare mapping efficiency (2), as HISAT2 showed relatively low mapping rates in some samples. The STAR aligner used the UCSC mm10 reference genome with a sjdbOverhang value of 250 bases.

**SAM/BAM Processing**

SAMtools was used to convert the resulting sequence alignment mapping (SAM) files to binary alignment mapping (BAM) format. Subsequently, the BAM files were sorted by genomic coordinates to facilitate downstream analysis (4).

**Transcript Assembly and Quantification**

StringTiewas used for transcript assembly and abundance estimation, using Ensembl reference annotation (release 91, Mus_musculus.GRCm38.91.gtf) (5). The following StringTie parameters were optimized for our experimental data:

- Minimum transcript length (-m): 100 bp
- Minimum junction coverage (-a): 5 reads
- Library type (-L): FR (forward-reverse) strand-specific
- Minimum read coverage (-s): 1.5
- Gap between read mappings (-g): 0

**Gene Expression Quantification**

After transcript assembly, HTSeq-count was used to quantify gene expression levels from aligned reads and assembled transcripts (1). The tool was configured with the following parameters:

- Input format (-f): BAM
- Strand specificity (-s): no (non-strand-specific library)
- Feature ID (-i): gene_id
- Feature type (-t): exon
- Mode (-m): intersection-strict

To enhance biological interpretation of results, additional attributes were extracted, including reference_id, ref_gene_id, ref_gene_name, and coverage (cov). The final gene count matrix was filtered to include only entries with validated mouse gene IDs (ENSMUS prefix), generating clean count files for downstream differential expression analysis.

**Single-cell RNA Sequencing Data Analysis**

This study used R language for single-cell RNA sequencing data analysis, primarily based on the Seurat package (7) for cell clustering and expression profile analysis.

**Data Integration and Preprocessing**

Gene IDs and expression values were extracted from count matrix files, with maximum expression values used to merge duplicate gene IDs. All sample data were integrated into a unified expression matrix, and missing values were replaced with 0.

**Quality Control and Normalization**

After constructing the Seurat object, the number of feature genes, total RNA counts, and mitochondrial gene percentage were calculated for each cell, and cell quality was assessed through scatter plots. Data was normalized using the LogNormalize method (scaling factor 10000), and the 2000 most variably expressed genes were selected for subsequent analysis.

**Dimensionality Reduction and Clustering Analysis**

Principal component analysis was performed on the normalized data, and UMAP and t-SNE dimensionality reduction were executed based on the first 20 principal components. Graph clustering algorithm (resolution=0.5) was used to identify cell subgroups, and clustering results were visualized through dimensionality reduction plots.

**Differential Expression and Marker Gene Identification**

The FindAllMarkers function was used to identify specific marker genes for each cluster (min.pct=0.01, logfc.threshold=0.1). The 10 or more most significantly differentially expressed genes in each cluster were selected to generate heatmaps, showing expression characteristics between clusters.

**Specific Gene Expression Analysis**

Expression patterns of key genes (*Tia1*, Kcnq1ot1, and *Gm26917*) were visualized on UMAP dimensionality reduction plots to explore the distribution characteristics of these genes in different cell subgroups.

**Single-cell Data Integration and Comparative Analysis**

This study conducted integrated analysis of two sets of single-cell RNA sequencing data (self-experimental data COSI and public dataset GSM3714747) to explore the heterogeneity of liver endothelial cells (10).

**GSM3714747 Dataset Processing**

Clustering results were imported from the preprocessed GSM3714747 dataset and reclusttered using a resolution parameter of 0.7. Based on gene expression characteristics, clustering results were relabeled as different anatomical region cell types in liver lobule, including PeriCentral1, PeriCentral2, PeriPortal, Core, PortalVein/HepaticArtery, CentralVein, and others.

Marker genes for each cell type were identified (min.pct=0.25, logfc.threshold=0.25), and the top 10 genes with the largest expression differences were selected to generate heatmaps.

**Data Integration Analysis**

Our COSI was integrated with the GSM3714747 (10) dataset, using FindIntegrationAnchors and IntegrateData functions. Standard procedures were applied to the integrated data for dimensionality reduction (PCA), clustering, and visualization (UMAP and t-SNE). Dimensionality reduction parameters were set to dims=1:20, min.dist=0.0, n.neighbors=200L, with a clustering resolution of 0.1.

**Subset Data In-depth Analysis**

Based on UMAP coordinates, the integrated data was filtered (UMAP1>-1.9) to extract cells from specific regions for in-depth analysis. The standard analysis workflow was re-executed on the subset data, with clustering resolution increased to 0.8 to reveal more refined cell subgroups. Differentially expressed genes were identified for each subgroup, and marker gene expression patterns were displayed through heatmaps.

**Experimental Sample Spatial Distribution Analysis**

For our COSI data, 384-well plate position information was analyzed by extracting row, column, and plate information from cell names and calculating the total gene expression for each sample.

**Morphology and Gene Expression Correlation Analysis**

**Data Processing and Visualization**

We constructed an R language-based analysis framework, integrating the Seurat package and various statistical analysis tools, to systematically evaluate single-cell morphological parameters. The analysis primarily focused on five key morphological characteristics: Mean Hole Area (Mean_Hole_Area), nuclear-to-cell ratio (Nuc2Cell), number of holes (Number_of_Holes), aspect ratio (AspectRatio), and eccentricity (Eccentricity).

**Feature Analysis Workflow**

Single Feature Comprehensive Analysis: Generated multi-dimensional views for each morphological parameter, including distribution plots, UMAP projections, cell type difference box plots, and gene correlation volcano plots. Distribution plots used density curves combined with histograms to show feature value distribution; UMAP projections displayed morphological feature distribution patterns in cell subgroup space; cell type difference analysis used box plots and statistical tests (t-test or ANOVA) to evaluate feature variation between different cell types;

gene correlation analysis screened genes significantly correlated with morphological features and presented them through volcano plots.

Inter-feature Comparison Analysis: Calculated correlation matrices between morphological parameters. Correlation strength between features was displayed through heatmaps.

Statistical Characteristic Evaluation: Calculated comprehensive statistical metrics for each morphological feature, including central tendency (mean, median), dispersion (standard deviation, IQR), distribution characteristics (skewness, kurtosis), etc. Data distribution characteristics were checked through histograms and box plots to evaluate the normality of morphological parameters.

Cell Population Difference Analysis: In-depth exploration of morphological feature differences among different cell types. ANOVA and Tukey HSD post-hoc tests were used for multiple group comparisons, and t-tests for two-group comparisons, with differences presented through box plots and bar charts.

**Redundancy Analysis (RDA) Revealing the Association Between Liver Sinusoidal Endothelial Cell Morphological Features and Gene Expression**

This study employed Redundancy Analysis (RDA) to explore the multivariate relationship between liver sinusoidal endothelial cell morphological features and gene expression patterns.

**Data Preparation and Preprocessing**

Morphological features included cell area (Area_Cell), nuclear area (Area_Nuc), nuclear-to-cell ratio (Nuc2Cell), eccentricity (Eccentricity), aspect ratio (AspectRatio), and hole-related features (Number_of_Holes, Mean_Hole_Area, etc.). After data cleaning and removal of samples with missing values, the morphological feature data was matched one-to-one with gene expression data.

**Redundancy Analysis Implementation**

The vegan package was used for redundancy analysis, with the transposed gene expression matrix as response variables and morphological features as constraint variables.

**Model Evaluation and Statistical Testing**

The RDA model was comprehensively evaluated: first, the overall model significance was assessed through permutation tests (anova.cca); then, independent conditional effect tests were conducted for each morphological feature to identify key morphological parameters significantly affecting gene expression. The proportion of constrained variation to total variation was calculated to quantify the degree to which morphological features explain gene expression variation.

#### Partial Least Squares Discriminant Analysis (PLS-DA)

##### Gene Expression Profile Analysis of Liver Sinusoidal Endothelial Cell Morphological Features

This study employed Partial Least Squares Discriminant Analysis (PLS-DA) to explore the association between liver sinusoidal endothelial cell morphological features and gene expression patterns.

##### Morphological Feature Stratification and Classification

For each morphological feature, cells were divided into three equal levels (low, medium, high) based on their distribution, creating categorical response variables.

##### PLS-DA Modeling and Cross-validation

For each morphological feature, a two-component PLS-DA model was built using the mixOmics package, with standardized processing to reduce the impact of dimensional differences. Model performance was evaluated through 5-fold cross-validation, calculating balanced error rates (BER) and the percentage of expression variation explained by each component to quantify the ability of different morphological features to distinguish cell types.

##### Loading Analysis and Feature Gene Identification

Loading values of the main components were extracted for each PLS-DA model, identifying the top 50 genes contributing most to morphological feature classification. The average expression levels of these genes were analyzed in cells with different morphological levels. For important genes, their loading values on the first and second principal components and their overall importance were further analyzed.

#### Gene Set Enrichment and Functional Analysis

##### Gene Set Enrichment Analysis and Correlation with Morphological Features

This study combined Gene Set Variation Analysis (GSVA) with correlation analysis to explore the association between liver sinusoidal endothelial cell morphological features and functional gene sets.

##### Data Preparation and Gene Set Acquisition

We first imported the integrated single-cell dataset and used the msigdbr package to obtain multi-level gene sets for mouse (Mus musculus) from the Molecular Signatures Database (MSigDB), including Hallmark pathways, cell type-specific gene sets, and tissue-specific gene sets. These gene sets were extracted as symbol identifiers and converted to the list format required for GSVA analysis.

**Gene Set Variation Analysis (GSVA)**

GSVA analysis was performed on single-cell transcriptome data to calculate activity scores for each gene set in each cell. Specific steps were as follows:

1. Extract gene expression matrix from Seurat object
2. Construct GSVA parameter object using gsvaParam and execute GSVA calculation
3. Extract target morphological feature values from metadata
4. Calculate Pearson correlation coefficients and *p*-values between GSVA scores for each gene set and morphological features

Correlation analyses were performed separately for five key morphological features (Mean_Hole_Area, Nuc2Cell, Number_of_Holes, AspectRatio, and Eccentricity), and results were integrated and stored.

**Correlation Analysis and Visualization**

Visualization and Analysis of Morphological Feature-Gene Set Relationships

The association between morphological features and gene set activity was assessed using multiple complementary approaches. Correlation clustering heatmaps were constructed for all morphological features and diverse gene sets. These heatmaps employed a blue-white-red chromatic gradient to represent negative to positive correlation values. Particular emphasis was placed on examining correlation patterns within Hallmark gene sets, liver-specific gene sets (AIZARANI_LIVER series), and kidney-related gene sets (KIDNEY series).

For each morphological feature, correlation-significance volcano plots were generated to visualize the statistical relationships with gene sets. These plots highlighted gene sets exhibiting significant correlations ($p<0.01$), with the top 30 most significantly correlated gene sets explicitly labeled, thereby facilitating identification of key functional pathways associated with specific morphological characteristics.

**Key Gene Set Screening and Extraction**

For each morphological feature, we defined a screening strategy based on statistical significance threshold ($p<0.01$). Through the extract_key_gene_sets function, up to 30 significantly correlated gene sets were extracted for each morphological feature. This layered screening approach ensured that we could capture the most relevant functional pathways for each morphological feature while maintaining interpretability and manageability of results.

**Keyword Frequency Analysis**

To reveal functional themes of gene sets, we constructed keyword extraction and statistical functionality. First, common prefixes (such as "GOBP_", "REACTOME_", "KEGG_", etc.) were removed through text preprocessing, then gene set names were tokenized, common connectors removed, and keyword frequency calculated. This method allowed us to identify core biological themes running through multiple gene sets, revealing patterns of association between morphological features and specific cellular functions.

#### Morphological Feature Analysis in Disease Models

##### Comparative Analysis of Liver Sinusoidal Endothelial Cell Morphological Features in Non-alcoholic Steatohepatitis Model (GSE129516, PRJNA531644)

This is a reanalysis of previously published data. This data set include cells from non-alcoholic steatohepatitis (NASH) model and control group (Chow)(10).

Evaluation of morphological feature differences between experimental groups was conducted using non-parametric statistical methodologies. A comprehensive three-tiered analytical approach was implemented:

For global difference assessment across all experimental groups, the Kruskal-Wallis rank sum test was employed. This non-parametric method was selected due to its robustness for multi-group comparisons with data that does not conform to normality assumptions.

Subsequent pairwise comparisons between individual groups were performed using the Wilcoxon rank sum test. To mitigate the elevated risk of Type I errors inherent in multiple comparison scenarios, the Bonferroni correction method was applied to adjust significance thresholds appropriately.

To examine broader disease-related differences, samples were stratified into two principal categories: Chow and NASH. The Wilcoxon test was subsequently utilized to evaluate significant morphological distinctions between disease state and control conditions, enabling assessment of pathology-associated structural alterations.

##### Temporal Change Analysis of Endothelial Cell Morphological Features in Diabetic Kidney Disease Model (GSE184652)

This is a reanalysis of previously published data. This data explored the effects of different therapeutic interventions on endothelial cell fenestrae structure and their temporal patterns by analyzing changes in morphological features of kidney endothelial cells in diabetic mice (9).

Data Preparation and Experimental Design

The experimental workflow was initiated with the importation of a kidney endothelial cell Seurat object previously processed with Gene Set Variation Analysis (GSVA). Subsequent regrouping of cells was performed in accordance with the experimental design parameters. The study incorporated multiple experimental groups derived from diabetic mouse models (db/db) subjected to diverse treatment regimens: the control group (db/m), untreated group (db/db), PBS control group, angiotensin-converting enzyme inhibitor (ACEi), rosiglitazone (Rosi), sodium-glucose cotransporter 2 inhibitor (SGLT2i), and combination therapy group. To facilitate assessment of temporal treatment effects, two distinct time points (2 days and 2 weeks) were established for each treatment regimen.

Temporal Analysis Methodology

Temporal alterations in morphological features were systematically evaluated through a two-tiered analytical approach:

Within-time point analysis was conducted by implementing Kruskal-Wallis rank sum tests independently on samples collected at 2 days (2d) and 2 weeks (2w) to determine global differences between treatment groups. Subsequent pairwise comparisons between individual treatment groups were performed using Wilcoxon tests, with Bonferroni correction applied to adjust for multiple comparisons.

Between-time points analysis was executed to assess the time dependency of treatment effects through systematic comparison of differences between identical treatment regimens at the 2-day and 2-week time points.

**Cell Culture and Staining Materials**

- 384-well plates (Aurora Microplates ABF101201A)
- RPMI 1640 medium (serum-free) (Invitrogen 11875-093)
- CellMask dye (1X and 3X concentrations) (Invitrogen C37608)
- PBS buffer
- 4% paraformaldehyde (PFA) (Electron Microscopy Sciences)
- DAPI dye (1X) (Invitrogen 62248)
- F-actin dye (1X) (Invitrogen A12380)
- Fluorescence microscope (Nikon)

**RNA Extraction and Library Construction Materials**

- Lysis buffer components:
    - 0.3% 2-mercaptoethanol (Sigma)
    - 0.1% Triton X-100 (Sigma)
    - 200 mM Tris-HCl (pH 8.0) (Sigma)
    - 1 mM $CaCl_2$ (Sigma)
    - Proteinase K (1:100 dilution) (Invitrogen 25530015)
- High-salt magnetic bead mixture:
    - 7% beads
    - 47% TCL buffer (Qiagen)
    - 47% 5M NaCl
- 80% ethanol
- Magnetic bead types:

- RNA Clean XP Beads (Beckman Coulter A63987)
  - DNA magnetic beads AMPure XP (Beckman Coulter A63881)
  - Magnetic separation rack for purification (Invitrogen)
- Reverse transcription reagents:
  - oligo dT-VM primer (0.06 μM)
  - dNTP mixture (NEB N0447S)
  - TS buffer (NEB M0466S)
  - TSO primer
  - TS enzyme (NEB M0466S)
  - $H_2O$ (nuclease-free water)
- PCR-related reagents:
  - KAPA HiFi PCR premix 2X (Roche 07958927001)
  - ISPCR primer (100 μM)
  - PCR thermal cycler
- Barcoding reagents (seqwell PW384-3B):
  - Sample barcode reagents (index set A: X021, X024, X038, X044)
  - Pooling barcode reagents
  - Encoding buffer
  - X termination solution
  - 10 mM Tris-HCl (pH 8.0)
- 384-well plates (for sample transfer)
- Electrophoresis system (for PCR product verification)

**Bioinformatics Analysis Software and Tools**

- FastQC: Sequencing quality control
- HISAT2 and STAR: Genome alignment tools
- SAMtools: SAM/BAM file processing
- StringTie: Transcript assembly and quantification
- HTSeq-count: Gene expression quantification
- R language and related packages:
  - Seurat: Single-cell RNA analysis
  - vegan: Redundancy analysis (RDA)
  - mixOmics: Partial least squares discriminant analysis (PLS-DA)
  - msigdbr: Gene set acquisition
  - GSVA: Gene set variation analysis
  - ggplot2 and ggpubr: Data visualization
  - lodash: Data processing
  - papaparse: CSV file processing

**Reference Chronic diseases Experimental Models (Literature Citations)**

The experimental models and data sources analyzed in this study are from the following published research. Our work involves analyzing these data rather than conducting original experiments:

- **Non-alcoholic steatohepatitis (NASH) model data** cited from public datasets (GSE129516, PRJNA531644), including:
  - Control group (Chow)
  - NASH group (special diet-induced)
- **Diabetic kidney disease model data (GSE184652)** from published literature, including the following experimental groups:
  - Control group (db/m)
  - Diabetic mice (db/db)
  - Various treatment group data:
    - PBS control group
    - ACEi (angiotensin-converting enzyme inhibitor) treatment group
    - Rosi (rosiglitazone) treatment group
    - SGLT2i (sodium-glucose cotransporter 2 inhibitor) treatment group
    - Combination therapy group
  - Sampling time points: 2 days (2d) and 2 weeks (2w)

The focus of this study is on computational analysis and integration of these published data, exploring the association between morphological features and gene expression patterns of liver sinusoidal endothelial cells and kidney endothelial cells, rather than conducting original animal experiments or disease model construction. We reanalyzed these data using advanced bioinformatics methods to gain new biological insights.

**Reference Data**

- Mouse reference genome (GRCm38)
- UCSC mm10 reference genome
- Ensembl reference annotation (release 91, Mus_musculus.GRCm38.91.gtf)
- Public datasets GSM3714747(10) and PRJNA531644 (GSE129516)(9)
- MSigDB Molecular Signatures Database:
  - Hallmark pathways
  - Cell type-specific gene sets
  - Tissue-specific gene sets (such as AIZARANI_LIVER series and KIDNEY series)
  - Other gene sets

**Statistical Analysis Tools**

- Kruskal-Wallis rank sum test
- Wilcoxon rank sum test
- Bonferroni correction
- Pearson correlation coefficient calculation
- Principal component analysis (PCA)
- *t*-test and ANOVA analysis
- Tukey HSD post-hoc test

**Supplementary Model**

This supplementary material describes a multi-target joint regression model designed for single-cell microscopy images, aimed at simultaneously predicting Number_of_Holes, Mean_Hole_Area, and Total_Hole_Area. Given the significant differences in imaging distributions between the *slide* (from Module 2) and *iSIM* (from Module 1) domains, the model does not learn directly from raw images; instead, it first incorporates an input domain alignment preprocessing module to uniformly correct the image intensity distribution of each sample. In the current optimal configuration, a global mean/standard deviation matching method—based on reference statistics derived from the *slide* training domain—is employed to map input images into the overall intensity space of the "*slide*" domain. This approach mitigates the impact of cross-domain pixel distribution shifts on downstream predictions, while the use of nan_to_num and numerical clipping ensures the numerical stability of the preprocessed images. The preprocessed images are then uniformly resized to 224×224 pixels to meet the requirements for batch training and input size consistency within the convolutional network. The network backbone utilizes a lightweight convolutional neural network, SmallCNN, comprising four sequential convolutional feature extraction modules. Each module consists of a convolutional layer, a batch normalization layer, a ReLU activation function, and a max-pooling layer, with channel counts progressively increasing in the sequence 3 → 32 → 64 → 128 → 256; this architecture facilitates the layer-by-layer extraction of features related to cell texture, boundaries, and internal structures. Following the final convolutional module, adaptive average pooling is applied to aggregate spatial features into a fixed-length global feature vector representation. This global feature vector is subsequently fed into a two-layer fully connected regression head, structured as: Flatten → Linear(256→128) → ReLU → Dropout(p=0.2) → Linear(128→3). The head ultimately outputs three continuous predicted values, corresponding to the three hole-related metrics. To mitigate discrepancies in scale and distribution among the three target variables, each target is individually subjected to Z-score standardization across the training set during the training phase; specifically, the target values are linearly normalized to a unified scale using the mean and standard deviation derived from the training set. Consequently, the model learns to predict these standardized regression values, which are then restored to their original physical scales via an inverse transformation during the inference phase. Comparative experiments indicate that, under the current small-sample conditions, Z-score standardization outperforms the log1p_zscore method. Furthermore, to address the issue of missing labels present in the raw data, this study performs label cleaning during the data loading phase: specifically, when Number_of_Holes = 0 and Total_Hole_Area = 0 while Mean_Hole_Area is missing, the sample is classified as a "no-hole sample" and the missing value is corrected to zero. Samples with completely missing labels are excluded during the training and supervised evaluation phases; however, they are retained during the unsupervised inference phase, where the model's predictions are outputted but do not contribute to the loss calculation. For model training, samples from the *slide* domain are utilized as training data, with a validation set partitioned from this data for model selection. The training employs SmoothL1Loss as the loss function and AdamW as the optimizer, with a learning rate set to $1 \times 10^{-4}$, supplemented by gradient clipping to enhance training stability. In the current experiments, the optimal configuration consists of global_mean_std preprocessing, slide_global_stats for reference statistics, a SmallCNN backbone, and z-score target normalization. The experimental results demonstrate that, following the completion of input domain alignment and label cleaning, the model training process no longer suffers from numerical explosion and is capable of generating finite and stable predictions on the *iSIM* dataset. The specific model architecture is shown in Figure S3. This architecture diagram was generated with the assistance of ChatGPT. The illustrations within the architecture diagram serve solely for conceptual demonstration and do not necessarily represent actual data.

### Figure S1

Comparison of Single Cell Images with Lowest and Highest Hole Counts

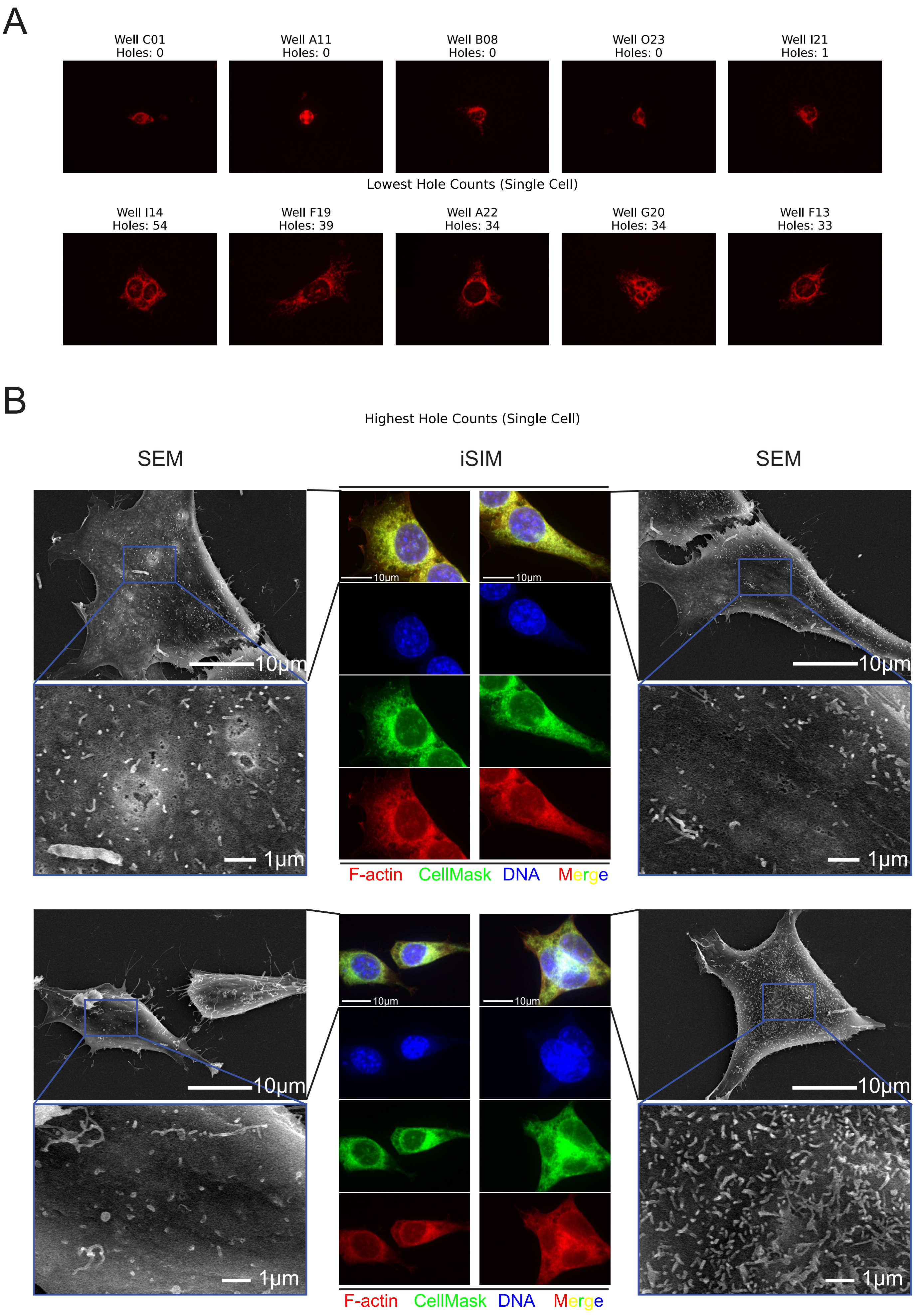

Figure S2

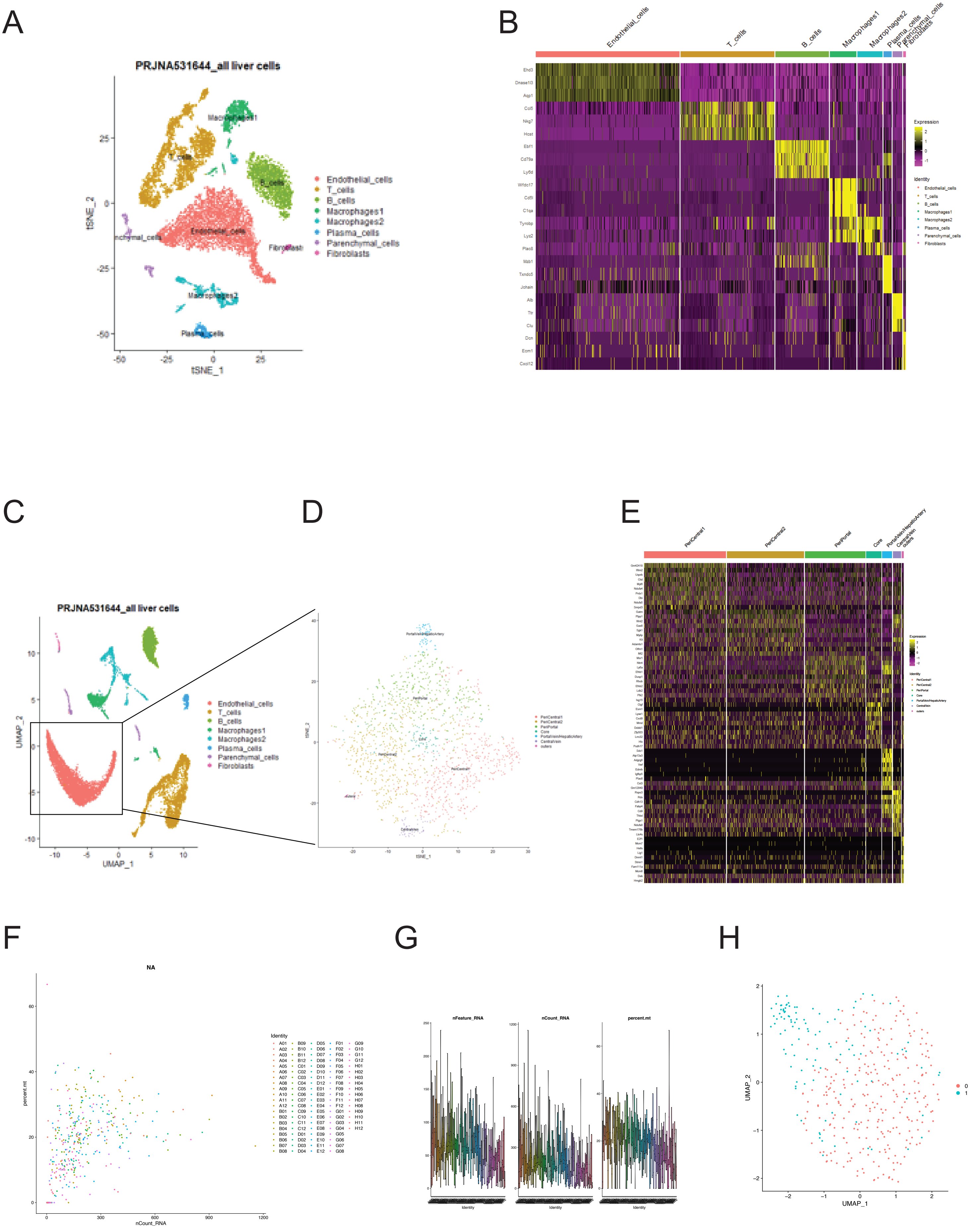
A
B
C
D
E
F
G
H
PRJNA531644_all liver cells
Endothelial_cells
T_cells
B_cells
Macrophages1
Macrophages2
Plasma_cells
Parenchymal_cells
Fibroblasts
tSNE_1
tSNE_2
UMAP_1
UMAP_2
Expression
Identity
PeriCentral1
PeriCentral2
PeriPortal
Core
PortalVein/HepaticArtery
CentralVein
outers
NA
percent.mt
nCount_RNA
nFeature_RNA
UMAP_1
UMAP_2

Figure S3

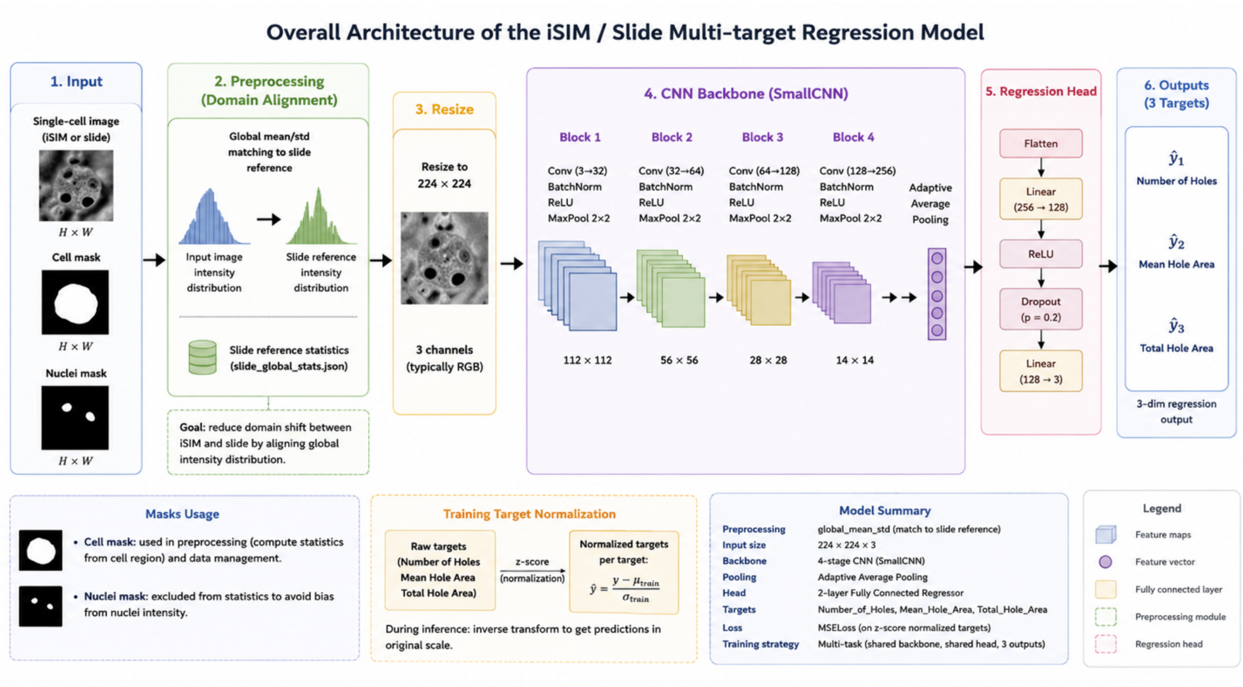

Overall Architecture of the iSIM / Slide Multi-target Regression Model
1. Input
Single-cell image (iSIM or slide)
H × W
Cell mask
H × W
Nuclei mask
H × W
2. Preprocessing (Domain Alignment)
Global mean/std matching to slide reference
Input image intensity distribution
Slide reference intensity distribution
Slide reference statistics (slide_global_stats.json)
Goal: reduce domain shift between iSIM and slide by aligning global intensity distribution.
3. Resize
Resize to 224 × 224
3 channels (typically RGB)
4. CNN Backbone (SmallCNN)
Block 1
Conv (3→32) BatchNorm ReLU MaxPool 2×2
Block 2
Conv (32→64) BatchNorm ReLU MaxPool 2×2
Block 3
Conv (64→128) BatchNorm ReLU MaxPool 2×2
Block 4
Conv (128→256) BatchNorm ReLU MaxPool 2×2
Adaptive Average Pooling
112 × 112
56 × 56
28 × 28
14 × 14
5. Regression Head
Flatten
Linear (256 → 128)
ReLU
Dropout (p = 0.2)
Linear (128 → 3)
6. Outputs (3 Targets)
ŷ1 Number of Holes
ŷ2 Mean Hole Area
ŷ3 Total Hole Area
3-dim regression output
Masks Usage
Cell mask: used in preprocessing (compute statistics from cell region) and data management.
Nuclei mask: excluded from statistics to avoid bias from nuclei intensity.
Training Target Normalization
Raw targets (Number of Holes Mean Hole Area Total Hole Area)
z-score (normalization)
Normalized targets per target:
$\hat{y} = \frac{y - \mu_{\text{train}}}{\sigma_{\text{train}}}$
During inference: inverse transform to get predictions in original scale.
Model Summary
Preprocessing global_mean_std (match to slide reference)
Input size 224 × 224 × 3
Backbone 4-stage CNN (SmallCNN)
Pooling Adaptive Average Pooling
Head 2-layer Fully Connected Regressor
Targets Number_of_Holes, Mean_Hole_Area, Total_Hole_Area
Loss MSELoss (on z-score normalized targets)
Training strategy Multi-task (shared backbone, shared head, 3 outputs)
Legend
Feature maps
Feature vector
Fully connected layer
Preprocessing module
Regression head
