## Supplementary figures and images for "Study on Liver Sinusoidal Endothelial Cell Fenestrations Based on Cellular Omics-Structure Integration Technology and Its Application in Metabolic Diseases"

### Supplementary file 29 Gene_Sets_volcano_plot.pdf

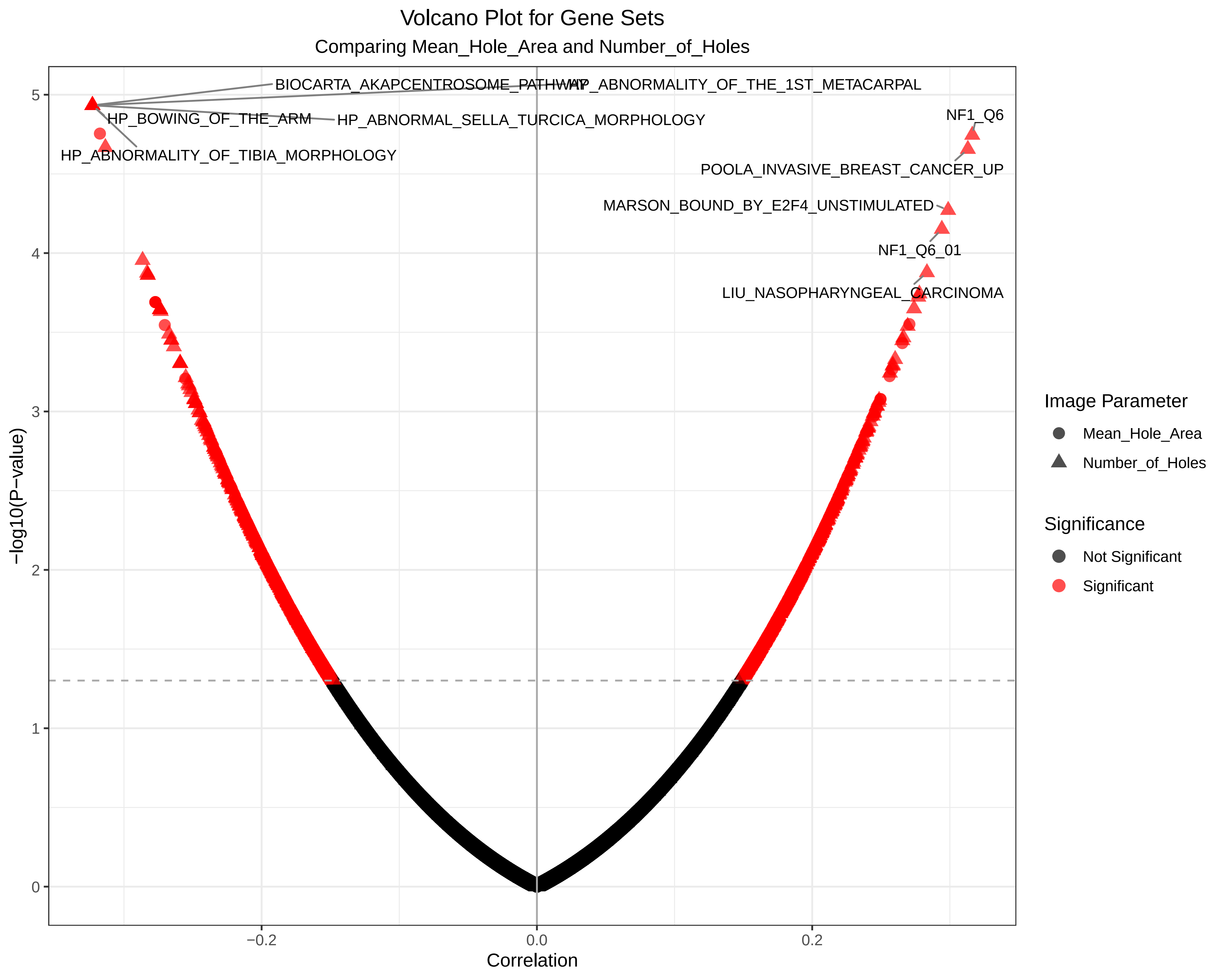

### Supplementary file 30 Mean_Hole_Area_volcano.pdf

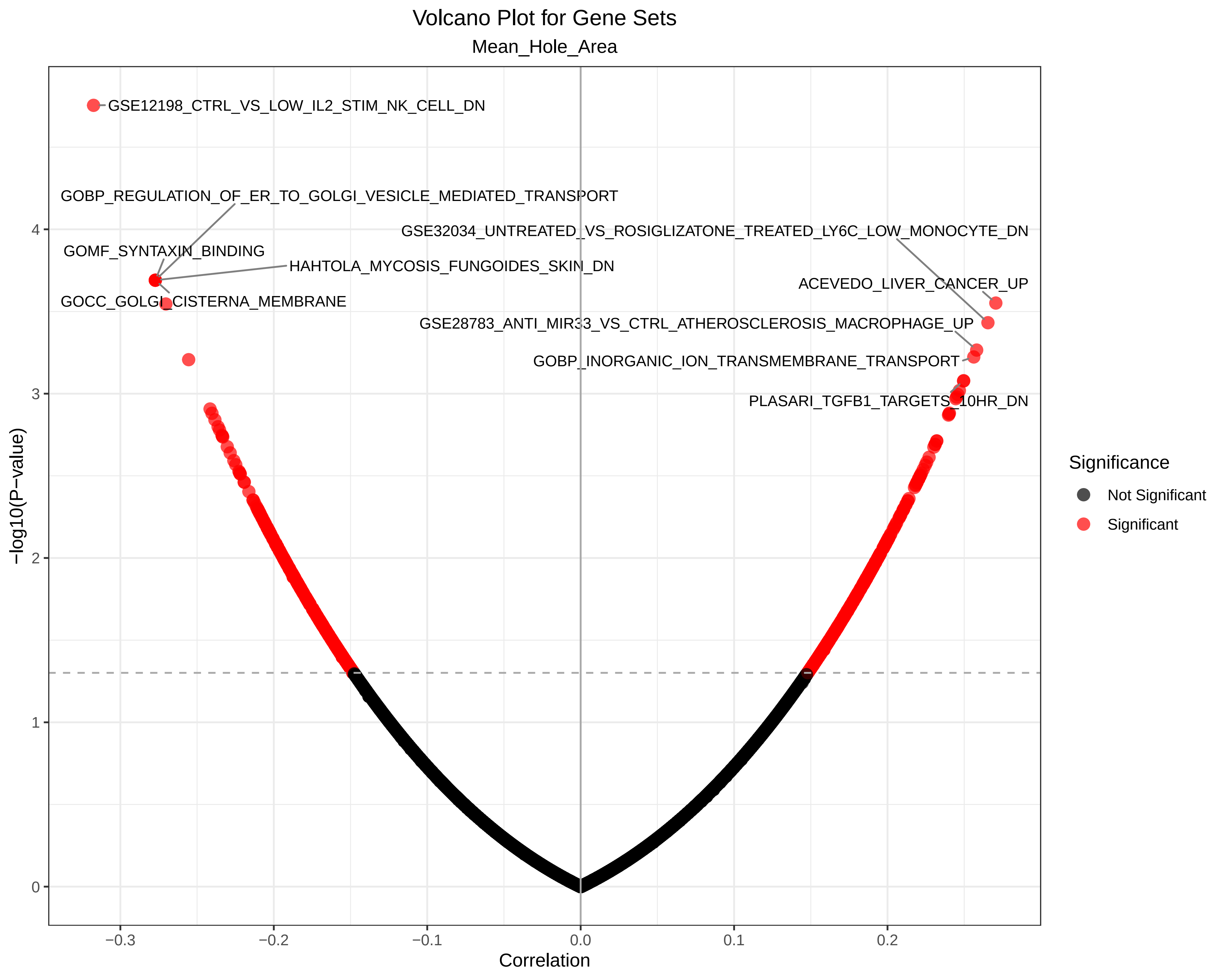

### Supplementary file 31 Number_of_Holes_volcano.pdf

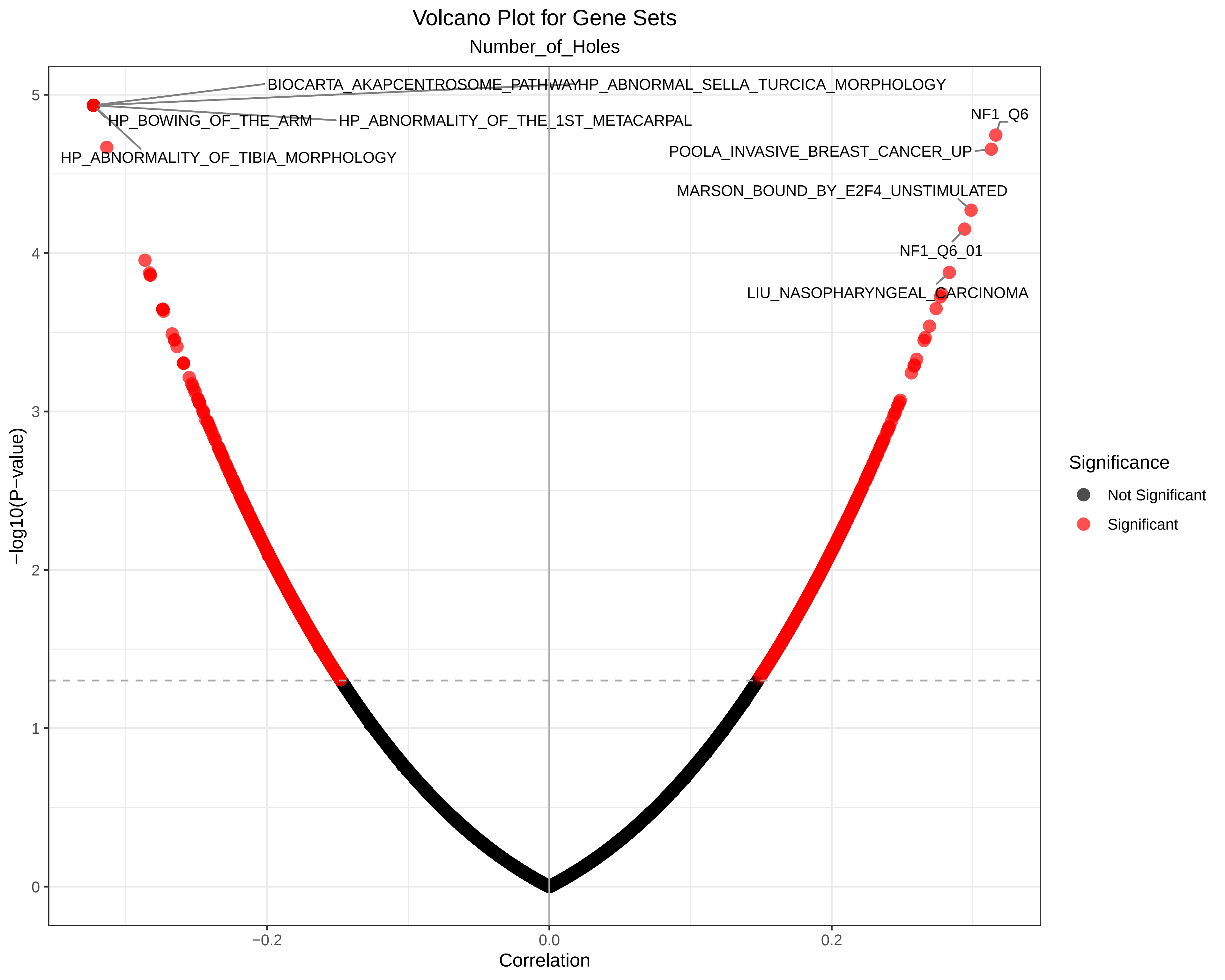

### Supplementary file 37 Correlation_Heatmap_Hallmark_Gene_Sets.pdf

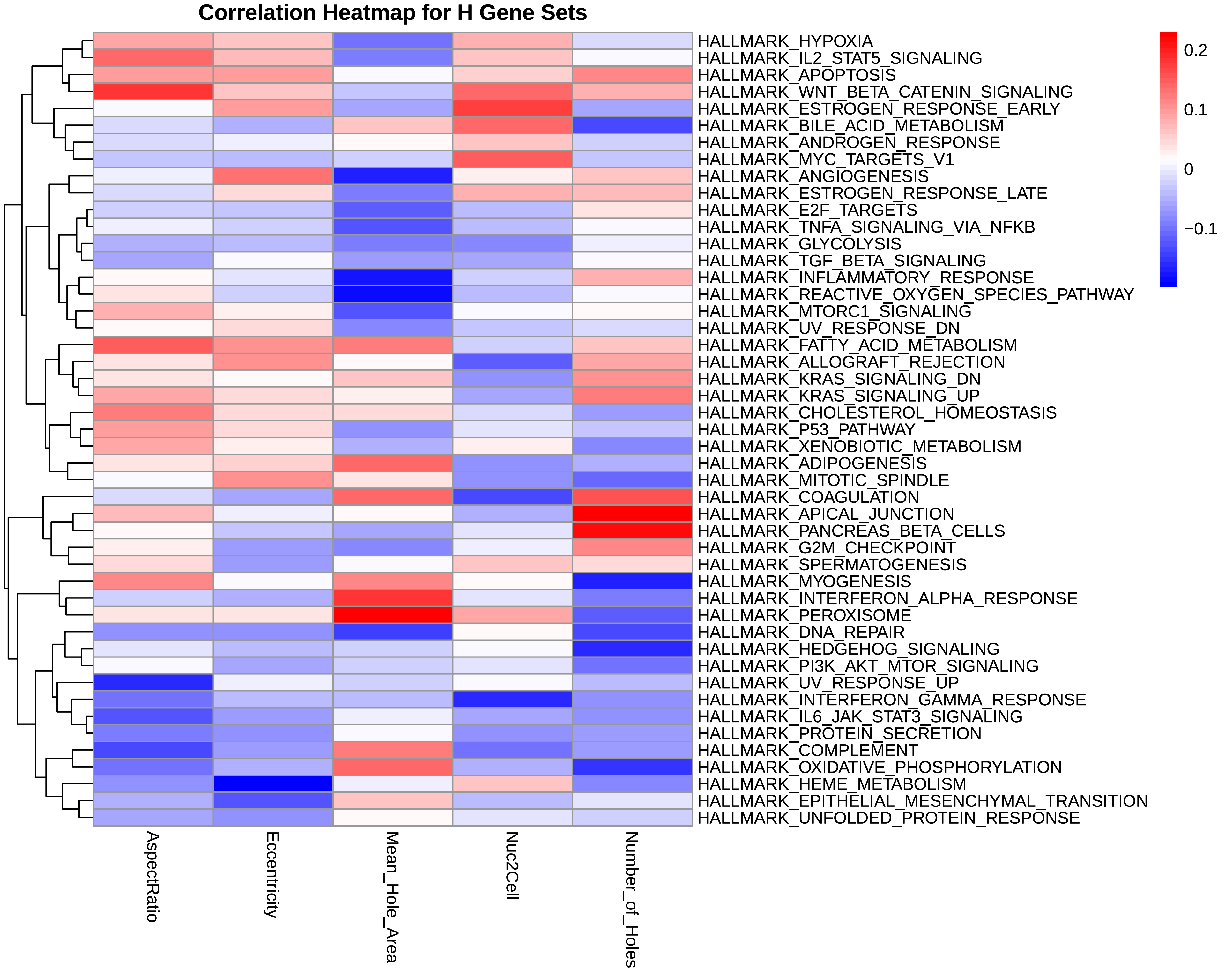

### Supplementary file 38 Correlation_Heatmap_KIDNEY_Gene_Sets.pdf

#### Correlation Heatmap for KIDNEY Gene Sets

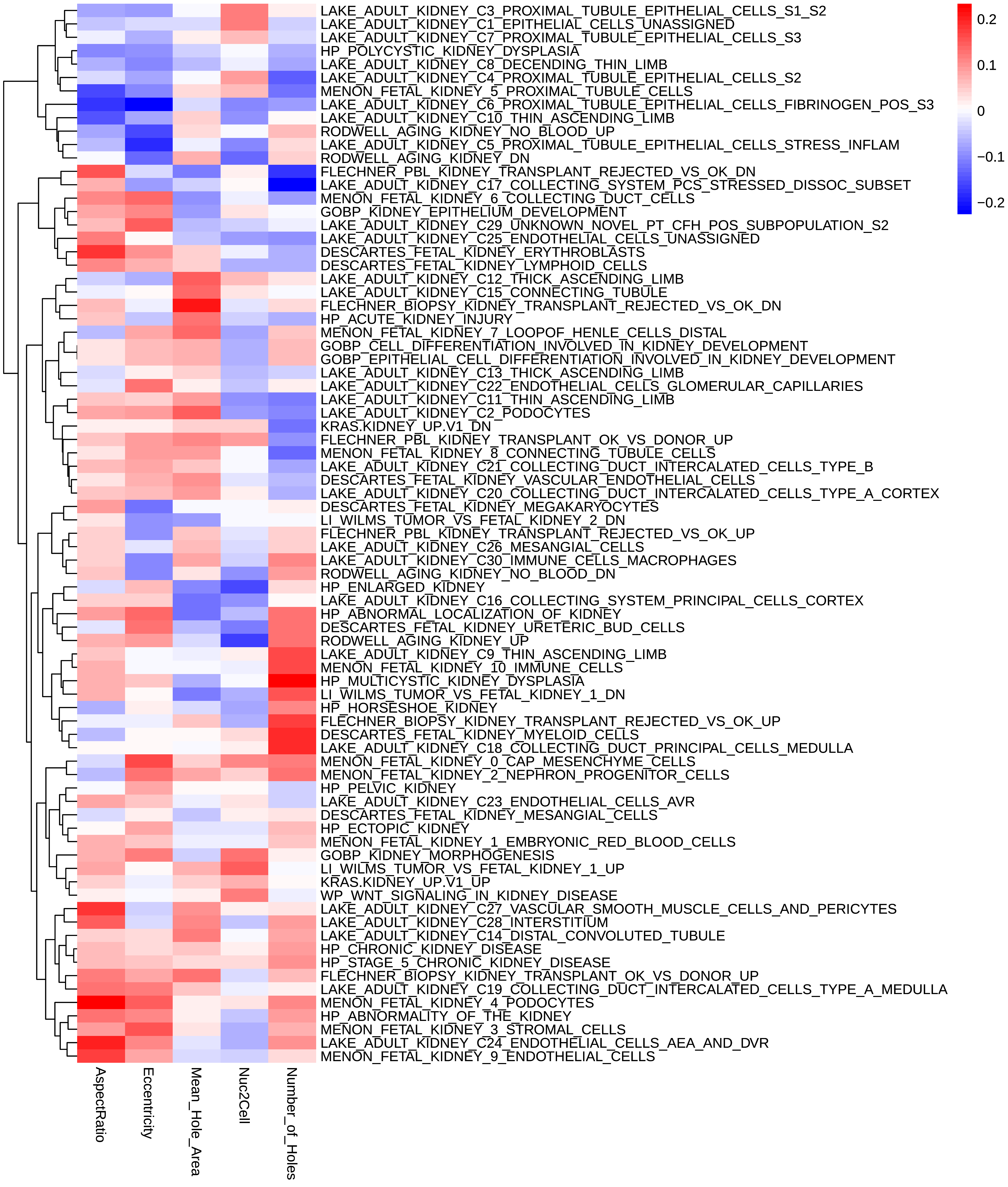

### Supplementary file 39 Correlation_Heatmap_C8_AIZARANI_LIVER_Gene_Sets.pdf

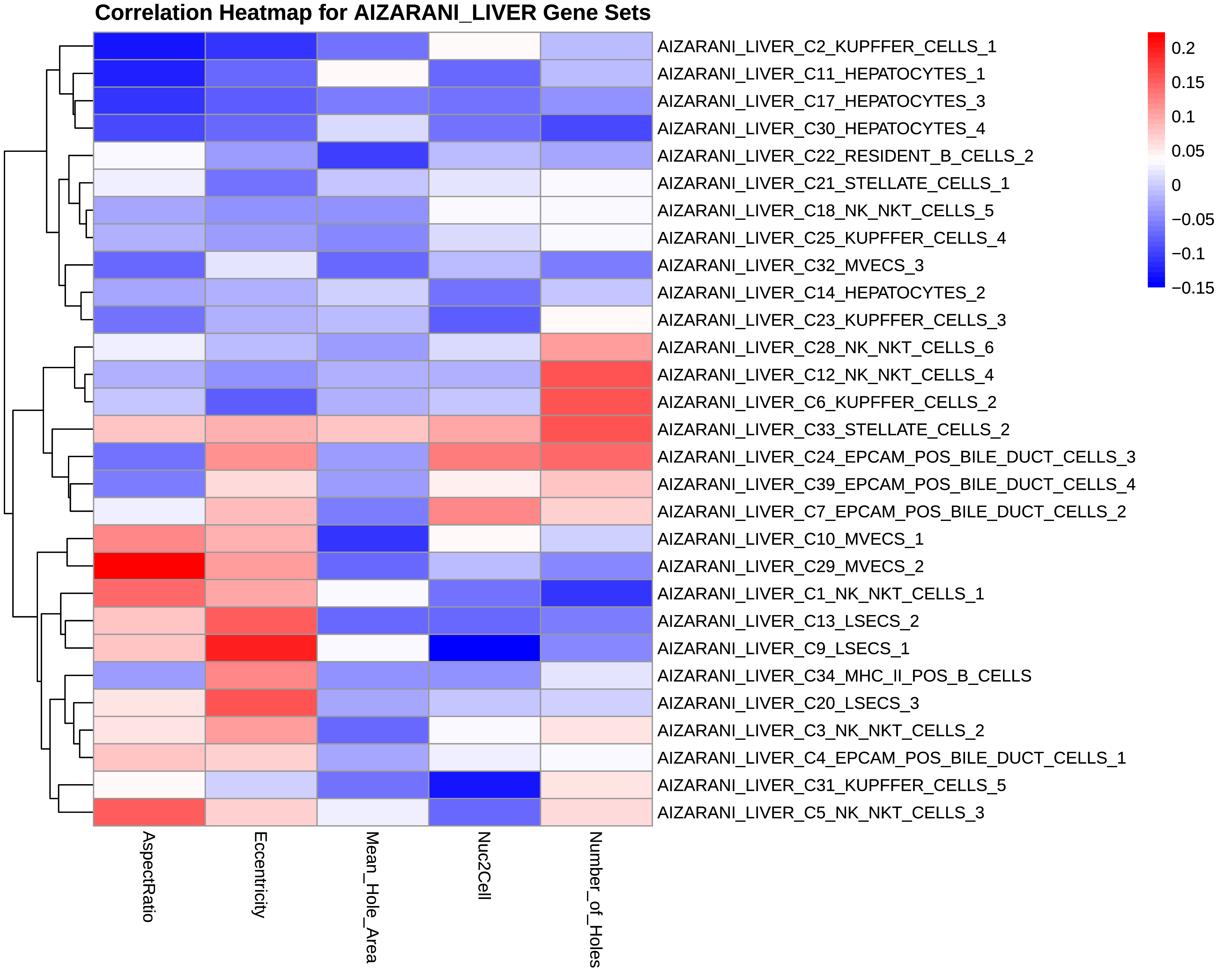
